# Activation of Toll and IMD pathways in the *Drosophila* brain following local and systemic bacterial infection

**DOI:** 10.1101/2025.10.20.683553

**Authors:** Sameekshya Mainali, Isaac Toles, Paige Magid, Jordan Grammer, Lauren Harper, Elizabeth Kitchens, Kaitlin Davis, Stanislava Chtarbanova

**Affiliations:** Department of Biological Sciences, University of Alabama, Tuscaloosa, AL-35487; Center for Convergent Bioscience and Medicine, University of Alabama, Tuscaloosa, AL-35487; Alabama Life Research Institute, University of Alabama, Tuscaloosa, AL-35487

**Keywords:** Brain infection, innate immunity, NF-κB, *Drosophila melanogaster*, antimicrobial peptides, glia

## Abstract

Brain infections are often life-threatening and have been linked to the development of neurodegenerative diseases. The fruit fly *Drosophila melanogaster* is a valuable experimental model to study immunity and the pathophysiology of brain infections. The exact cellular pathways through which brain-specific immune responses are mounted in *Drosophila*, however, remain poorly characterized. Here, we investigated how brain-specific or systemic infection with *Micrococcus luteus* and *Escherichia coli* bacteria activates the *Drosophila* NF-κB innate immune pathways Toll and immune deficiency (IMD) in the central nervous system of the fly. We tested the hypothesis that these pathways are acutely activated in the *Drosophila* brain, and that their activation persists over time, even if bacteria have been cleared. We demonstrate that in control genotypes, brain-specific bacterial infection leads to *Drosomycin* (*Drs*, Toll pathway) and *Diptericin B* (*DiptB*, IMD pathway) upregulation and that glia appear to be the primary cell type mounting this immune response at both early and later stages of infection, although some activation is observed in neurons as well. We show that the upregulation of *Drs* and *DiptB* expression also depends on canonical components of the Toll and IMD pathways, respectively. Interestingly, we found that systemic infection with *M. luteus* leads to brain-specific *Drs* activation and that signals from the fat body and hemocytes can activate the Toll pathway in the brain, pointing to an inter-organ communication. Together, these results contribute to our understanding of how non-lethal bacterial infections result in activation of NF-κB immunity in *Drosophila* brain that could potentially be targeted to prevent progression of neurodegeneration.

**Highlights:**

- Brain immunity is induced following bacterial brain infection and depends on canonical NF-κB pathway components.
- NF-κB signaling pathways are induced acutely and persist over time after bacterial brain infection.
- Host functional immunity clears bacteria in the brain post-bacterial brain infection.
- Glia are the main brain cell type in which NF-κB immunity is induced at both early and later stages of bacterial infection.

## Introduction

Central nervous system (CNS) infections caused by bacteria, viruses, fungi, or parasites can lead to long-term neurological damage or death. The human CNS is protected by an innate immune system primarily via resident immune cells such as microglia and astrocytes. These cells are crucial for mounting a defense against various external threats, including infections and injuries (Lye and Chtarbanova, 2018; Rodríguez et al., 2022). Following a brain infection, an innate immune inflammatory cascade is initiated, which proceeds through multiple cellular and molecular phases that help in clearing the pathogen (Lotz et al., 2021; Zindler and Zipp, 2010). This immediate, short-term acute immune response is beneficial and promotes tissue healing (Postolache et al., 2020). However, the nature of this immune response can shift from protective to detrimental. For instance, in the context of brain injuries, prolonged and chronic activation of the immune system, particularly the sustained response of macrophages (including microglia), can lead to a secondary phase of tissue damage. This chronic neuroinflammation is a significant factor in long-term functional loss. It is characterized by the persistent release of inflammatory molecules that can be toxic to neurons and other surrounding brain cells (Larrea et al., 2023; Swanson et al., 2020a; Winkler et al., 2025). Thus, appropriate immune reactions in the brain are indispensable to reducing the harmful effects of infection and maintaining overall brain health.

The fruit fly *Drosophila melanogaster* is a powerful model organism that shares conserved Nuclear Factor kappa B (NF-κB) innate immune signaling pathways with mammals, including the Toll pathway, which is similar to mammalian Toll-like receptor (TLR) and Interleukin-1 receptor (IL-1R) pathway, and the immune deficiency (IMD) pathway which is similar to the mammalian Tumor Necrosis Factor Receptor (TNFR) pathway. Following systemic infection, activation of both pathways depends on the recognition of microbial cell wall components or virulence factors by *Drosophila* pattern recognition receptors (PRRs) (Lemaitre and Hoffmann, 2007; Liegeois and Ferrandon, 2022). The Toll pathway is activated by microbial peptidoglycans through activation of an extracellular ligand, Spatzle (Spz) (Leclerc and Reichhart, 2004). The Peptidoglycan recognition protein-SA (PGRP-SA)/Gram-negative bacteria binding protein 1 (GNBP1) complex, which is necessary for recognizing and responding to Gram-positive bacteria or the Gram-negative bacteria binding protein 3 (GNBP3), essential for sensing fungal infections, initiates a signaling cascade that leads to cleavage of Spz involving multiple proteolytic activities, particularly Spatzle-Processing Enzyme (SPE) (Jang et al., 2006; Shan et al., 2023). Spz is synthesized as an inactive precursor molecule, which is cleaved into its active form by SPE. Cleaved Spz, binds to the Toll receptor, which is located on the cell membrane. This recognition initiates a downstream signaling pathway that involves phosphorylation and degradation of Cactus, an inhibitor of the NF-κB transcription factors Dorsal and Dorsal-related immunity factor (Dif) (Lemaitre and Hoffmann, 2007; Rutschmann et al., 2000). Upon Cactus degradation, Dif translocates to the nucleus and initiates transcription of many genes, including genes encoding antimicrobial peptides (AMPs), with *Drosomycin* being a key target (Lemaitre et al., 1997). On the other hand, DAP-type peptidoglycan of Gram-negative bacteria binds to the transmembrane receptor PGRP-LC that is found in the cell membrane, or the intracellular receptor Peptidoglycan Recognition Protein-LE (PGRP-LE), and activates the IMD pathway (Liegeois and Ferrandon, 2022). Upon binding, a cascade of signaling events occurs, recruiting a complex formed of the adaptor proteins Imd, Fas-associated death domain (Fadd), and a caspase, Death-related ced-3 (Dredd). This complex activates the downstream IMD signaling components, resulting in the activation of the NF-κB transcription factor Relish (Rel). Rel translocates into the nucleus and initiates the transcription of many genes, including genes encoding AMPs, with *Diptericin* being a key target (Myllymaki et al., 2014). Subsequently, these AMPs directly target pathogens and combat microbial threats.

To prevent overstimulation and possible tissue damage, immune activation in *Drosophila* is also kept in check via negative regulatory mechanisms (Lee and Ferrandon, 2011). This is particularly important in tissues such as the nervous system, which have minimal regenerative capacity. In *Drosophila*, mutations in several negative regulators resulting in excessive NF-κB pathway activation lead to neurodegeneration, altered locomotor behavior, and shorter lifespan (Cao et al., 2013; Kounatidis et al., 2017). Additionally, direct introduction of a bacterial mix of *Micrococcus luteus* (*M. luteus*, Gram-positive bacteria) and *Escherichia coli* (*E. coli*, Gram-negative bacteria) into the fly brain leads to an NF-κB-mediated inflammatory response, which causes progressive neurodegeneration accompanied by impaired locomotor behavior (Cao et al., 2013). Understanding the upstream steps of innate immune activation in the brain could therefore provide new insights about the pathophysiology of neurodegeneration.

While systemic activation of innate immunity in response to bacterial pathogens in *Drosophila* has been characterized in great detail, the activation and the nature of innate immune reactions in tissues such as the brain remain understudied. One study showed that systemic infection performed by injecting *Drosophila* larvae with Group B Streptococcus (GBS) bacteria (Gram-positive bacteria) can trigger an inflammatory response via the activation of NF-κB signaling, which also leads to the recruitment of hemocytes (the fly equivalent to mammalian macrophages) to the brain (Winkler et al., 2021). Because GBS is lethal to larvae it is difficult to examine neuroinflammatory responses that may persist following bacterial challenge and contribute to development of neurological sequalae over time. To address this gap in research, in this study, we utilized adult *Drosophila melanogaster* to study infection-induced neuroinflammation.

We investigated how non-lethal brain infection and systemic infection with *M. luteus* and *E. coli* affect immune activation in the brain of control genotypes and various mutants of the Toll and IMD pathways. We show that bacterial infection leads to an acute (6-48h) *Diptericin B* (*DiptB)* and *Drosomycin* (*Drs*) upregulation in *wild-type* (WT) control brains, and that this activation persists for up to two weeks following the initial challenge. We find that the IMD pathway mediates *DiptB* expression in the brain following *E. coli* brain infection and that the Toll pathway mediates *Drs* expression in the brain following *M. luteus* brain infection. We further show that functional immunity cleared the bacterial load in the brain 12h post-brain infection indicating that the long-term activation of immunity is not associated with the presence of live bacteria that have not been cleared by the host. We also demonstrate that systemic *M. luteus* infection leads to *Drs* expression in the brain of control genotypes, activation that is mediated via signals from the periphery. Using immunohistochemistry and a co-localization approach, we also show that following bacterial challenge, Toll and IMD pathways are primarily activated in glial cells in the brain.

## Materials and methods

### Drosophila stocks and handling

*Drosophila* stocks were amplified and maintained on standard Nutri-Fly® Bloomington formulation food (Cat #: 66-113) in a 25°C incubator. 0-3 days-old flies were collected and aged for 2-3 days before experiments were conducted. For experiments involving aging up to two weeks old, flies were flipped every 2-3 days in a fresh food-containing vial until the desired age was reached. The following *Drosophila* stocks were used as control genotypes: *y^1^ w^67C23^* (BL_6599), *Dipt-LacZ, Drs-GFP, y^1^, w\** (BL_55707), *w^1118^* (gift from Dr. R. John Manak), and *Relish*^+/+^ (genetic background control for *Relish^del^*mutants; a gift from Dr. David Wassarman; (Swanson et al., 2020b)). *AMP::GFP* flies including *Attacin::GFP* and *Cecropin::GFP* were obtained from Dr. David Wassarman, originally described in (Tzou et al., 2000). RNAi lines used include UAS-*MyD88*^RNAi^ (BL_36107) and UAS-*Spz*^RNAi^ (BL_34699). GAL4 driver lines used include *ppl-GAL4* (BL_58768), *Hml-GAL4* (BL_30140), *Repo-GAL4* (BL_7415). Mutants used in this study include *Dipt-LacZ, Drs-GFP, y^1^, w*, cn^1^, bw^1^, Dif^1^* (BL_36559), *Tak1^2^*(BL_26272), *Dredd^B118^* (BL_55712), *Myd88* mutants (*MyD88^c03881^*allele (Tauszig-Delamasure et al., 2002), kind gift from Dr. Dominique Ferrandon), *Relish^E20^* (BL_55714), and *Relish^del^*(gift from Dr. David Wassarman, (Swanson et al., 2020b)). Only male flies were used in all experiments.

### Bacterial strains and infections

The stock bacterial cultures of Gram-positive *Micrococcus luteus* (Ward’s Science #85WOGG6), Gram-negative *Escherichia coli* (ATCC #11775), and Ampicillin-resistant *Escherichia coli* (HB101 + pGLO) were obtained from the Microbiological collection at the Department of Biological Sciences at The University of Alabama. Liquid bacterial cultures were grown in an Erlenmeyer flask containing 50 mL of LB media (*M. luteus* and *E. coli*). Ampicillin-resistant *E.coli* were grown in an Erlenmeyer flask containing 50 mL of LB media supplemented with 100 ug/mL Ampicillin (Alfa Aesar, Cat # J60977). Flasks were shaken overnight in the shaker at 161 rpm at 30°C for *M. luteus* and 37°C for *E.coli.* The next day, the optical density of the overnight suspension was measured, adjusted to OD_600nm_ = 0.2, and centrifuged at 4000 rpm for 10 minutes at 4°C. After centrifugation, the culture supernatant was discarded, and the obtained pellets were used for the infection experiment. To prepare heat-killed bacterial cultures, the overnight-grown bacterial suspensions were autoclaved at liquid cycle for 30 min at 121°C and 15psi. Autoclaved bacterial suspensions were streaked on Tryptic Soy Agar (TSA) plates to verify that the heat treatment worked (no bacterial colonies were observed after 48h incubation at 37°C for *E. coli* and 30°C for *M. luteus*). Non-infected flies were directly processed for experiments without any injection treatment. For sterile injury and bacterial infection, flies were subjected to the following treatments-Sterile injury: Flies were directly pricked either into the head through the left or right eye (brain infection) or the cuticle on one side of the thorax (systemic infection) with a thin minutien pin (0.1 mm) (Roboz Surgical, Cat. # RS-6083-10) mounted on a pin holder. Before the poke, the minutien pin was briefly dipped into EtOH70% solution and gently wiped with a Kim wipe. Bacterial infection: The minutien pin was dipped into the concentrated live or heat-killed bacterial pellets between each fly. Like the sterile injury, flies were injected either in the head or the thorax (brain and systemic infection, respectively). Following injection flies were incubated at 25°C until used for experiments.

### Bacterial load quantification

For bacterial load quantification in *Drosophila* heads, overnight-grown Ampicillin-resistant *Escherichia coli* (HB101 + pGLO) and *Micrococcus luteus* (Ward’s Science #85WOGG6) were used to cause brain and systemic infection, as described above. Brain- and systemically infected flies with either of the bacteria were incubated at 25°C for 12 hours and 48 hours post-infection to calculate colony forming units (CFUs) per 5 heads. For bacterial load determination in whole flies, flies systemically infected with ampicillin-resistant *E. coli* were incubated at 25°C for 12 hours and 24 hours post-infection to calculate CFUs per whole fly. After incubation at respective time points, either 5 heads or a whole fly were transferred into a microcentrifuge tube containing 100 µL of sterile PBS1X and homogenized for ∼3-4 min using a sterile pestle. After homogenization, 1:100 serial dilutions were performed, and 10 µL of the 10^5, 10^6, and 10^7 dilutions from the homogenates were plated on TSA plates and incubated at 37 °C for 24 hours for *E. coli* infected samples and at 30 °C for 48 hours for *M. luteus* infected samples. Bacterial colonies were counted, and the number of colonies per mL of a sample was calculated by using the formula: CFU/mL (number of colonies * dilution factor)/ volume transferred onto the plate.

### RNA extraction and gene expression analysis

*Drosophila* brains were dissected using forceps (Dumostar style 5, biological forceps, Electron Microscopy Sciences, Cat. # 72705-01) wiped with EtOH70%, by holding the sharply pointed forceps in the fly’s head and tearing apart the cuticle from the head. Brains were dissected at the rate of ∼ 40-50 brains/hour. ∼15 dissected brains were gently transferred into a microcentrifuge tube containing 400 uL of an RNAlater solution (Thermo Fischer, Cat. # AM7020) and stored at −20°C until further used. RNA was extracted from ∼15 brains using the Quick-RNA™ Micro Prep Kit (Zymo Research, Cat. # R1050) according to the manufacturer’s instructions, with the following modification: 1. Brains were homogenized for 2 minutes instead of the recommended 1 minute. 2. The DNAse1 treatment was extended from 20 min to 30 min at room temperature and centrifuged for 1 min instead of 30 sec. 3. After adding 700μL of RNA Wash Buffer to the filter column, centrifugation was done for 1 min instead of 30 sec. 4. For the final RNA elution step, 15μL of DNase/RNase-Free Water was directly added to the column and centrifuged for 1 min instead of 30 sec. The concentration (ng/μL) of eluted RNA was measured using the NanoDrop One (Thermo Scientific™). The extracted RNA was stored at −20°C in the freezer until it was used for cDNA synthesis. The High-Capacity cDNA Reverse Transcription kit (Applied Biosystems, Cat. #: 4368814) was used to synthesize cDNA according to the manufacturer’s protocol. The initial RNA mass used for all samples was 250 ng. The reaction cycle was: 10 minutes at 25°C, 120 minutes at 37°C, 5 min at 85°C and pause at 4°C. Samples were stored at −20°C until further use. For the real-time qPCR reaction, cDNA was diluted 5 times. All samples were run in technical triplicates and nuclease-free water was used as a negative control. Briefly, each reaction consisted of 5 *μ*L Power Track Sybr Green (Applied Biosystems), 0.5 *μ*L forward primer (10*μ*M), 0.5 *μ*L reverse primer (10*μ*M) and 3 *μ*L nuclease-free water that was added to 1 *μ*L of diluted cDNA. The housekeeping gene *RpL32* was used as an endogenous control for the normalization of gene expression. The genes of interest measured in this study include *Diptericin B* (*DiptB*) and *Drosomycin* (*Drs*). Sequences of Forward (FW) and Reverse (RV) primers are as follows (in 5’è 3’ orientation): *RpL32*: FW: AAGAAGCGCACCAAGCACTTCATC, RV: TCTGTTGTCGATACCCTTGGGCTT; *Drs*: FW: CGTGAGAACCTTTTCCAATATGATG, RV: TCCCAGGACCACCAGCAT (Cao et al., 2013); *DiptB*: FW: ACCGCACTACCCACTCAAT, RV: GGTCCACACCTTCTGGTGAC (Cao et al., 2013); *BomS2*: FW: AGTCGTCACCGTCTTTGTGTT, RV: CAGTATTTGCAGTCCCCGTTG (Xu et al., 2023). The qPCR reaction was carried out in a Step-One-Plus machine (Applied Biosystems) using the standard two-hour quantitative analysis including a melt curve reaction. The following reaction cycle conditions were: 95°C for 10 minutes, 40 repetitions of 95°C for 15 seconds followed by lowering the temperature to 60°C for 1 minute. Then, one round of 95°C for 15 seconds, 60°C for 1 minute, and finally 95°C for 15 seconds. Relative gene expression was determined using the formula 2^ (*RpL32* C_t_ value) / 2^ (the gene of interest C_t_ value) and was log 2-transformed prior statistical analyses.

### Immunostaining and imaging

5-day-old male *Attacin::GFP*, *Cecropin::GFP*, and *Dipt-LacZ, Drs-GFP, y^1^, w\** flies were subjected to different injection treatment conditions. Brains were dissected at different time points post-infection: 24 hours (to address early immune activation) and 2 weeks (to address long-term immune activation). Brains were dissected in PBS1X and were fixed using Paraformaldehyde (4%PFA) in PBS1X for 30 min by gently rotating on a shaker. Next, the fixative solution was removed, and the brains were washed with 0.1% PBS-Triton X (PBS-T) 3 times on a shaker, each for at least 15 minutes at room temperature. After the third wash, brains were incubated in a blocking solution using PBS-T with 4% Normal Goat Serum (MP Biomedicals, Cat. # ICN19135680) for 1 hour at room temperature. Following this step, primary antibodies diluted in blocking solution were added to the brains and incubated overnight in the refrigerator at 4°C. The primary antibodies were used as follows: chicken-anti-GFP (1:1000) (Invitrogen, Cat. # A10262), Mouse-anti-Repo (1:50) (DSHB, Cat. # 8D12), the monoclonal antibody developed by Corey Goodman was obtained from the Developmental Studies Hybridoma Bank, created by the NICHD of the NIH and maintained at The University of Iowa, Department of Biology, Iowa City, IA 52242, Rat-anti-Elav (1:100) (DSHB, Cat. # 7E8A10), the monoclonal antibody developed by Gerald M. Rubin was obtained from the Developmental Studies Hybridoma Bank, created by the NICHD of the NIH and maintained at The University of Iowa, Department of Biology, Iowa City, IA 52242, and Rabbit-anti-ß-galactosidase (1:100) (Invitrogen, Cat. # A11132). After removing primary antibodies, brains were washed with PBS-T on a rotating shaker 3 times for 15 minutes at room temperature. Then, secondary antibodies diluted in 0.1% PBS-T were added to the brains, and samples were covered with aluminum foil to protect them from light and incubated for 3 hours at room temperature on a rotating shaker. The secondary antibodies were used as follows: goat-anti-chicken Alexa Fluor 488 (1:500) (Invitrogen, Cat. # A11039), goat-anti-mouse Alexa Fluor 568 (1:500) (Invitrogen, Cat. # A11031), goat-anti-rat Alexa Fluor 633 (1:500) (Invitrogen, Cat. # A21094), goat-anti-rabbit Alexa Fluor 405 (1:1000) (Invitrogen, Cat. # A31556). After removing secondary antibodies, brains were washed with PBS-T on a rotating shaker 3 times for 15 minutes at room temperature. Finally, brains were mounted on a microscopic slide into a drop of ProLong™ Diamond Antifade Mountant (Invitrogen, Cat. # P36961) and covered with a glass coverslip. Z-stacks of entire brains were obtained under a 20X objective on a confocal microscope (Nikon Eclipse Ti2 Laser Scanning Confocal Microscope) using the same camera settings for all treatment conditions and time points. To detect specific morphological characteristics of glial cell subtypes, the 40X oil immersion objective was used to obtain higher-magnification images. Acquired images at both 20X and 40X were further analyzed using Fiji ImageJ2 (Version: 2.3.0/1.53q) software. Unless otherwise specified, a maximum projection of 3 slices per sample from n=3 brains were examined per experimental condition. Images were enhanced using the ‘Brightness and contrast’ function in Fiji ImageJ2 to improve visualization; however, measurements of fluorescence intensity were done on unmanipulated files. Fluorescence intensity plots (GFP, Repo and Elav) were generated by selecting a region of interest (ROI) using a single image chosen from the corresponding z-stacks. Measurements were done using the ‘Plot profile’ function in Fiji ImageJ2 for the same ROI across all fluorescence channels.

### Statistical analysis

Statistical analysis was done using GraphPad Prism v.10 software. Statistics are based on a 2-way ANOVA test, with Tukey’s post-test for multiple comparisons. P-values of P < 0.05 were considered significant.

## Results

### Canonical IMD pathway components mediate DiptB expression in the brain following *E. coli brain infection*

To test whether IMD pathway components are implicated in response to *E. coli* infection, we tested *Tak1^2^* and *Dredd^B118^* mutants in the IMD pathway. We measured *Diptericin* (*DiptB*) 6 h post-injury (hpi) in brain samples of the control genotype *y^1^w^67C23^*and *Tak1^2^* and *Dredd^B118^* flies after brain and systemic injection, respectively. At 6 hpi we observed that in brains of the control strain both sterile injury and *E. coli* infection resulted in significant *DiptB* upregulation (P=0.0062 and P<0.0001, respectively) compared with non-injected flies **(Fig. 1A, Table S1)**. However, the difference between sterile brain injury and *E. coli* brain infection was not significant (P=0.2359), suggesting that at this time point brain injury alone resulted in upregulation of *DiptB*. When compared to systemic *E. coli* infection, brain *E. coli* infection resulted in significantly higher *DiptB* brain expression (P=0.0003) **(Fig. 1A, Table S1)**. Similarly, we did not observe a significant difference in *DiptB* expression in the brain of the control strain between sterile brain injury and sterile systemic injury (P=0.2299) **(Fig. 1A, Table S1)**. As additional controls activating the immune system without causing live infection, we performed brain and systemic injections using heat-killed *E. coli* (HK *E. coli*). We observed significant *DiptB* induction in the brain of the control strain post-HK *E. coli* brain infection compared to non-infected cohorts (P<0.0001) **(Fig. 1A, Table S1)**. However, we did not observe significant *DiptB* induction in the brain of the control strain post-HK *E. coli* brain infection compared to sterile brain injured cohorts (P=0.4131) **(Fig. 1A, Table S1)**. Moreover, we observed significant *DiptB* induction in the brain of the control strain post-HK *E. coli* brain infection in comparison to post-HK *E. coli* systemic infection (P=0.0006) **(Fig. 1A)**. Systemic infection did not significantly induce *DiptB* in the brain of the control strain post*-E. coli* when compared to systemic sterile injury (P>0.9999) **(Fig. 1A, Table S1)**. Both IMD pathway mutants *Tak1^2^* and *Dredd^B118^* failed to induce *DiptB* expression after both brain and systemic live and HK *E.coli* infection, suggesting that *DiptB* expression in the brain depends on these canonical components of the pathway **(Fig. 1A, Table S1)**. Altogether, these results indicate that brain-specific, but not systemic infection with live or HK *E.coli* stimulates the upregulation of *DiptB* in the brain, and that this response is IMD pathway-dependent.

**Figure 1.**
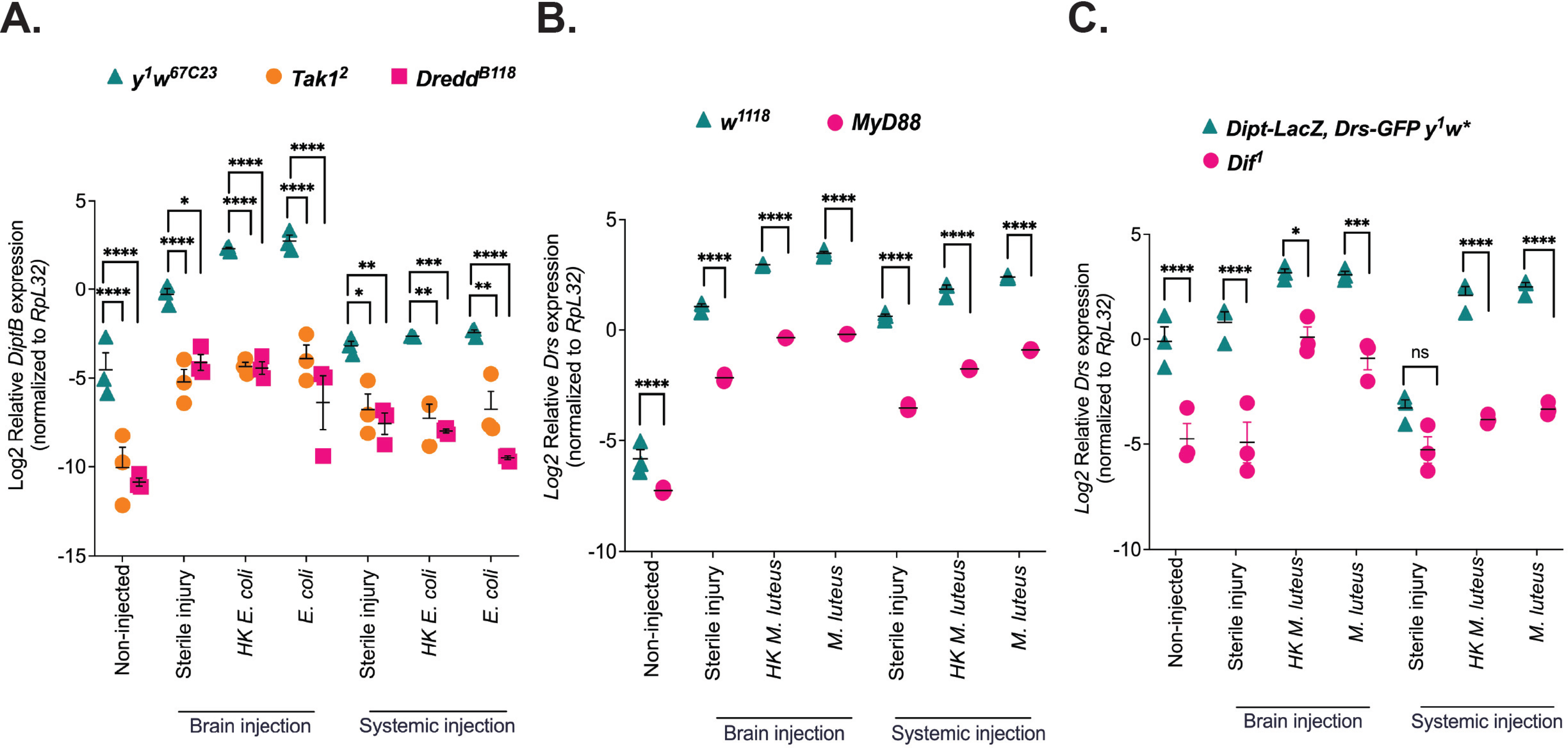
Dependence of brain-specific AMP expression on *Drosophila* IMD and Toll pathways. **(A)** *DiptB* gene expression was measured in control genotype and IMD pathway mutants’ brain tissue 6h post-head (brain) and thorax (systemic) *E. coli* infection. **(B-C)** *Drs* gene expression was measured in control genotypes and Toll pathway mutants’ brain tissue 24h post-head (brain) and thorax (systemic) *M. luteus* infection. N =15 dissected brains were used per replicate, with n=3 biological replicates (individual symbols on the graph) for each experimental treatment. **(A-C)**, *DiptB* and *Drs* expression levels were normalized to the housekeeping gene *RpL32* and log transformed. Mean ± SEM are shown and asterisks shown above the standard deviation bars denote significant differences in the pairwise control comparisons. *: P<0.05, **: P<0.01, ***: P < 0.001, ****: P<0.0001, ns: not significant based on a 2-way ANOVA test, with Tukey’s post-test for multiple comparisons.

### Canonical Toll pathway components mediate Drs gene expression in the brain following M. luteus brain infection

We next evaluated the contribution of the Toll pathway in the activation of brain immunity following infection with the Gram-positive bacterium *M. luteus*. We assayed brain expression of the AMP gene *Drosomycin* (*Drs*) in two Toll pathway mutants (*MyD88* and *Dif^1^*) and their controls (*w^1118^* and *Dipt-LacZ, Drs-GFP y^1^w\**, respectively) at 24 hpi. **(Fig. 1B, C, Table S1).** We observed that following sterile brain injury in comparison to the non-infected cohort, *Drs* gene expression was significantly induced in the *w^1118^* brain (P<0.0001), but this was not the case in *Dipt-LacZ, Drs-GFP y^1^w\** brains (P=0.9888) **(Fig. 1B, C, Table S1)**. This could represent a genotype-specific difference in response to brain injury. However, in both control strains, brain-specific *M. luteus* infection resulted in significantly induced *Drs* expression in comparison to non-infected cohorts (P<0.0001 for *w^1118^*, **Fig. 1B**, and P=0.0089 for *Dipt-LacZ, Drs-GFP y^1^w\**, **Fig. 1C, Table S1)**. *Drs* was significantly induced in the *w^1118^* brains post-*M. luteus* brain infection in comparison to post-sterile brain injury (P<0.0001) **(Fig. 1B, Table S1)**. However, this difference was not significant in the brains of the second control genotype (P=0.1575) **(Fig. 1C, Table S1)**. We didn’t observe significant *Drs* expression in *w^1118^* brains post-sterile brain injury compared to sterile systemic injury (P-value=0.5878) **(Fig. 1B, Table S1)**. However, the second control genotype strain *Dipt-LacZ, Drs-GFP y^1^w\** displayed a significant difference in brain *Drs* expression between the sterile brain injury cohort compared to the sterile systemic injury cohort (P=0.0004) **(Fig. 1C, Table S1)**. HK *M. luteus* brain injection resulted in significant *Drs* induction in comparison to sterile brain injured cohorts (P<0.0001) and non-injected cohorts (P<0.0001) **(Fig. 1B)**. *Drs* induction was significant in *w^1118^* brains post-HK *M. luteus* brain infection in comparison to post-HK *M. luteus* systemic infection (P=0.0003) **(Fig. 1B, Table S1)**. However, this difference in *Drs* expression was not significant in the second control genotype (P=0.9622) **(Fig. 1C, Table S1)**. *Drs* expression was significantly lower in the two Toll pathway mutants we tested (*Myd88* and *Dif)* after brain and systemic live and HK *M. luteus* infection. As an additional readout of Toll pathway activation, we measured *BomS2* gene expression in *w^1118^* and *MyD88* post-brain and systemic *M. luteus* infection. *BomS2* expression was significantly lower in *MyD88* mutant compared to *w^1118^* post brain *M. luteus* infection (P=0.0129), and systemic *M. luteus* infection (P=0.009), suggesting that *Drs* and *BomS2* upregulation following infection in the brain is Toll pathway-dependent **(Fig. 1B, 1C, and Fig. S1) (Table 1)**. Interestingly, we didn’t observe significant *Drs* expression in the *w^1118^* brain post systemic vs brain *M. luteus* infection (P= 0.9532) **(Fig. 1B, Table S1)**. Similarly, we didn’t observe significant *Drs* expression in *Dipt-LacZ, Drs-GFP y^1^w\** brain post systemic vs brain *M. luteus* infection (P= 0.9999) **(Fig. 1C, Table S1)**. Together, these results indicate that both brain-specific and systemic infection with *M. luteus* trigger *Drs* upregulation in the brain, which also appears to depend on the Toll pathway.

### Signals from the fat body and hemocytes activate the Toll pathway in the brain, inducing Drs expression

We next investigated how systemic *M. luteus* infection leads to brain *Drs* expression **(Fig. 1B, C, Table S2)**. The hypothetical model **(Fig. 2A)** outlines the idea that following systemic infection, the upstream component of the Toll pathway *Spz*, a circulating cytokine in the hemolymph (HL), which is synthesized mainly by fat body and hemocytes, can in its active form following processing by Spatzle Processing Enzyme (SPE) diffuse and reach the brain to stimulate Toll pathway activation in this tissue. We hypothesized that Spatzle (Spz) is synthesized by organs outside the brain following *M. luteus* systemic infection and released in the hemolymph. After activation, Spatzle would circulate and bind the Toll receptor on different tissues, including the fat body and hemocytes in the periphery, as well as glial cells in the nervous system, activating the Toll pathway, inducing brain-specific *Drs* activation. To test this hypothesis, we knocked down the genes encoding *Spz* in the fat body (*Ppl-Gal4>UAS-Spz RNAi*), hemocytes (*Hml-Gal4>UAS-Spz RNAi*), and glia (*Repo-Gal4>UAS-Spz RNAi*) **(Fig. 2 A-C, Table S2)** and knocked down the adaptor *MyD88*, which functions downstream of the Toll receptor in the fat body (*Ppl-Gal4>UAS-MyD88 RNAi*), hemocytes (*Hml-Gal4>UAS-MyD88 RNAi*), and glia (*Repo-Gal4>UAS-MyD88 RNAi*) **(Fig. 2 D-F, Table S2)** and measured *Drs* expression 24h post-brain and systemic *M. luteus* infection. Flies expressing one copy of each of the *Gal4* drivers used served as controls. In the case of direct brain infection, in driver controls, we observed significant *Drs* expression at 24h post-brain *M. luteus* infection compared to non-injected cohorts (P <0.0001, **Fig. 2 B-G, Table S2)**. Similarly, *Drs* expression in the brain was significantly upregulated in driver controls post-brain *M. luteus* infection compared to their sterile brain-injured counterparts (P <0.0001, **Fig. 2 B-G, Table S2)**. These results recapitulate the upregulation of *Drs* in two control genotype brains following *M. luteus* infection of this tissue (**Fig. 1 B, C, Table S2**). While in both *Spz* and *MyD88* KD flies, *Drs* expression was significantly upregulated post-brain *M. luteus* infection compared to their non-injected and sterile brain-injured counterparts (P<0.0001, **Fig. 2 B-G, Table S2)**, the expression of this AMP was significantly reduced in comparison to their respective driver controls (P<0.0001, **Fig. 2B-G, Table S2)**. In the case of systemic infection, in driver controls, we observed significantly increased *Drs* brain expression at 24h post-systemic *M. luteus* infection compared to non-injected and sterile systemic-injured cohorts (P<0.0001, **Fig. 2 B-G, Table S2)**. This is consistent with the upregulation of *Drs* following systemic *M. luteus* infection in the brains of the two control genotypes (**Fig. 1 B, C, Table S2**). In both *Spz* and *MyD88* KD flies, *Drs* expression was significantly upregulated post-systemic *M. luteus* infection compared to the non-injected and sterile brain-injured cohorts (P<0.0001, **Fig. 2 B-G, Table S2)**. However, compared to their respective driver controls, both *Spz* and *MyD88* KD flies displayed significantly reduced *Drs* expression in the brain post-systemic *M. luteus* infection (P<0.0001, **Fig. 2 B-G, Table S2)** (P<0.0001). This suggests that Spz, which is required to activate the Toll pathway in the brain, could be produced by each of those tissues — i.e., both systemic (fat body/hemocytes) and locally in the brain (glia). Independent of MyD88, as long as more Spz is produced by the fat body and hemocytes, it presumably gets into the brain and activates the Toll pathway, inducing *Drs* expression. Since Spz is an upstream component of Toll signaling we decided to knock down *MyD88* and see if the downstream adaptor knockdown reduces response in the receiving tissue and possibly fits the canonical pathway order (Toll → MyD88 → Tube → Pelle → Dif/Dorsal → target AMPs like *Drosomycin*). Our results imply that both systemic (fat body and hemocyte) and local brain (glia) cells receive MyD88/Toll signals, resulting in intracellular pathway activation required for brain *Drosomycin* induction. Altogether, these results suggest that the brain’s response to systemic infection follows the canonical Spz →Toll →MyD88 signaling in brain *Drs* induction following *M. luteus* infection.

**Figure 2.**
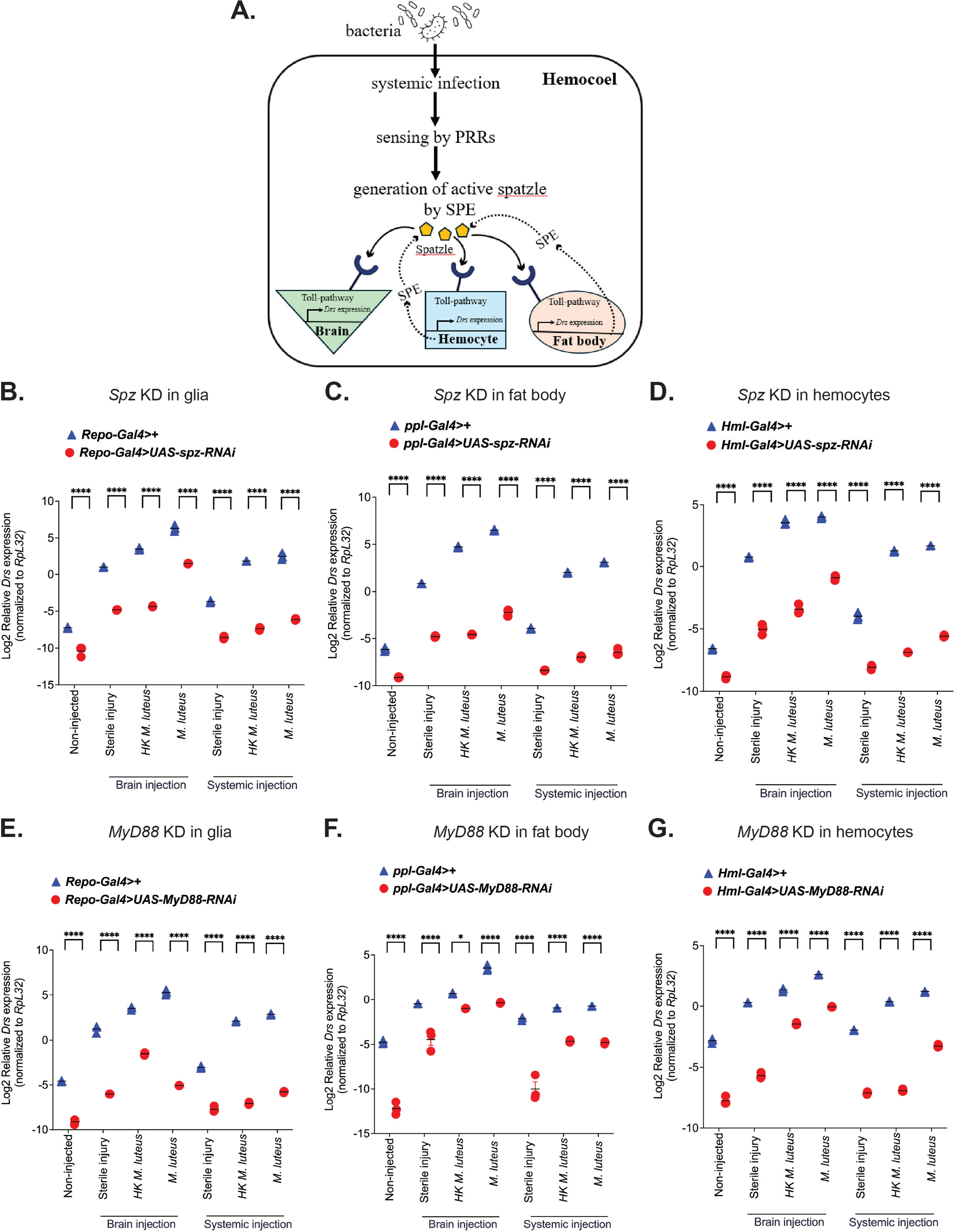
Fat body, hemocytes, and glia contribute to brain *Drs* upregulation following systemic *M. luteus* infection. **(A)** Proposed working model showing systemically derived Spz activating the Toll pathway in the glia following *M. luteus* infection. **(B-D)** *Drs* gene expression was measured in *Spz* knockdown flies and their respective Gal-4 driver controls at 24h post-brain and systemic *M. luteus* infection. *Spz* knockdown was performed in glia, fat body, and hemocytes, respectively. **(E-G)** *Drs* gene expression was measured in *MyD88* knockdown flies and their respective Gal-4 driver controls at 24h post-brain and systemic *M. luteus* infection. *MyD88* knockdown was performed in glia, fat body, and hemocytes, respectively. *Drs* expression levels were normalized to the housekeeping gene *RpL32* and log transformed. Mean ± SEM; ***P < .005 based on a 2-way ANOVA test, with Tukey’s post-test for multiple comparisons. ns, not significant. Asterisks shown above the standard deviation bars denote significant differences in the pairwise control comparisons.

### DiptB and Drs expression is activated in the brain acutely after bacterial brain infection, and this activation persists over time

To further assess acute and long-term immune activation of the IMD and Toll pathways in response to direct brain infection with *E. coli* and *M. luteus*, we measured *DiptB* and *Drs* gene expression at subsequent time points: 6-48h and 2 weeks. In *y^1^ w^67C23^* (control strain used for IMD pathway mutants) *E. coli* infected brains, at 6 hpi, we did not observe a significant *DiptB* expression compared to sterile injured cohorts (P>0.9999). However, at 12h, 24h, 36h, 48h, and up to 2 weeks (P<0.0001), the effect of the injury resolved **(Fig. 3A, Table S3)**. Moreover, *DiptB* expression in *E. coli-*infected brains at 6 hpi. was not significantly different from *DiptB* expression in *E.coli-*infected brains for the remaining time points, such as 12h (P=0.3276), 24h (P=0.9278), 36h (P=0.8148) and 48h (0.3708), except 6h vs 12h, which is highly significant (P<0.0001) and persisted for up to 2 weeks, but not significant (P=0.9694) **(Fig. 3A, Table S3)**. Similarly, in *w^1118^* and *Dipt-LacZ, Drs-GFP y^1^w\** (two control strains used for Toll pathway mutants) *M. luteus* infected brains, at 24 hpi, *Drs* was significantly induced in *M. luteus*-infected brains compared to sterile-injured cohorts (P=0.0015) **(Fig. 3B, Table S3**) and (P=0.0029) **(Fig. 3C, Table S3)**. At 36h post-*M. luteus* infection, *Drs* was significantly upregulated in *w^1118^* control brains compared to 24 hpi (P= 0.0019), and it persisted for up to 2 weeks, although *Drs* expression is non-significant compared to 24h post *M. luteus* infection (P=0.0734) **(Fig. 3B, Table S3)**. In *Dipt-LacZ, Drs-GFP y^1^w\** control brains, *Drs* persistently activated up to 2 weeks, although the *Drs* expression is not significant compared to 24h post *M. luteus* infection (P=0.4371) **(Fig. 3C, Table S3)**. Taken together, these results show that *DiptB* is upregulated acutely from 6-48h post- *E. coli* brain infection, and that *Drs* is upregulated from 12h to 48h post- *M. luteus* brain infection. In comparison to sterile-injured cohorts, the expression of both genes remained significantly upregulated for up to 2 weeks post-brain bacterial infection, indicating long-term activation of the immune response.

**Figure 3.**
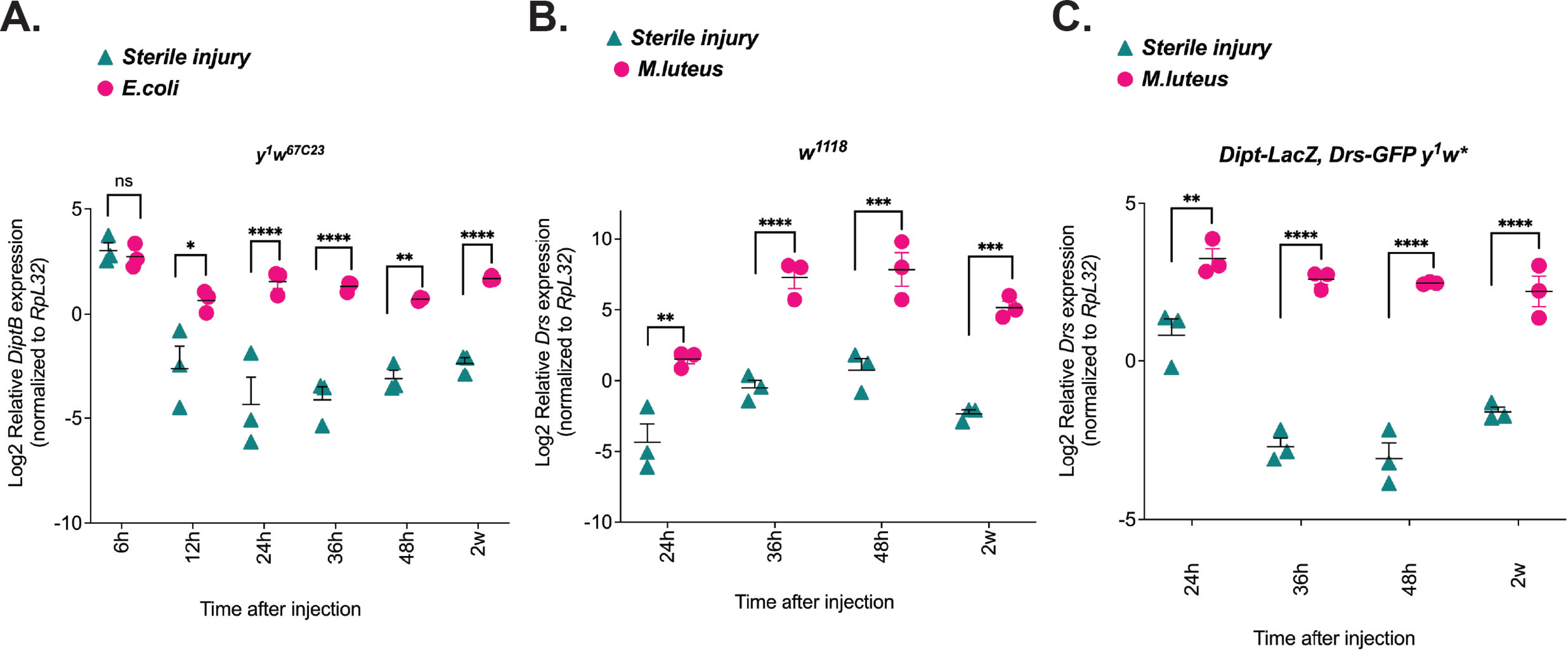
NF-κB brain immunity is acutely induced following direct bacterial challenge. The immune response persists over time, indicating long-term immune activation. **(A)** *DiptB* gene expression was measured in control *y^1^ w^67c23^*genotype brains at different time points (6h-48h) and 2 weeks post *E. coli* brain infection to assay acute and long-term immune activation, respectively. **(B-C)** *Drs* gene expression was measured in *w^1118^*and *Dipt-LacZ, Drs-GFP y^1^w\** control genotype brains at different time points (24h-48h) and 2 weeks post *M. luteus* infection to assay acute and long-term immune activation, respectively. Both *DiptB* and *Drs* expression levels were normalized to the housekeeping gene *RpL32*. Mean ± SEM; ***P < .005 based on a 2-way ANOVA test, with Tukey’s post-test for multiple comparisons. ns, not significant. Asterisks shown above the standard deviation bars denote significant differences in the pairwise control comparisons.

### Host functional immunity clears the bacterial load in control flies but not in NF-kB mutants

Previously, in control genotype flies, we observed that host immunity was activated for up to 2 weeks after bacterial challenge raising the possibility that bacteria persist and stimulate immune responses in the brain. Therefore, to see if this long-term activated immunity is the result of non-cleared bacteria in the brain, we counted bacterial colonies in whole heads post-brain and systemic infection by performing a CFU assay. At 0h post-brain and systemic *E. coli* infection, we did not observe a significant difference in bacterial colony count in the head of WT control flies and mutants, except for *Rel^+/+^* vs *Rel^E20^*(P=0.0354) **(Fig. 4A, Table S4, S5)**. At 12h post-brain and systemic infection with *E. coli* and *M. luteus*, bacteria were completely cleared from the head of control strains in all experiments **(Fig. 4A-D, Table S4, S5).** At 12h post-brain and systemic *E. coli* infection, bacterial colonies in the mutants’ head (*Rel^del^* mutant and *Rel^E20^*) (P< 0.0001, **Fig. 4A-B, Table S4, S5)** persisted at significantly higher levels compared to *Rel^+/+^*. At 48h, *E.coli* was cleared in the *Rel^+/+^*head, whereas mutants died post-brain and systemic *E. coli* infection. The systemic *E. coli* infection in the whole fly was used as an additional control to verify that *Rel^del^* mutants accumulated more CFUs until they died at 48h post-infection **(Fig. S2, Table S4, S5)**. Similarly, at 0h post-brain and systemic *M. luteus* infection, we didn’t observe a significant difference in bacterial colony count in the heads of *Dipt-LacZ, Drs-GFP y^1^w\** controls, and *Dif^1^* mutant post-brain (P=0.7116) **(Fig. 4C, Table S4, S5)** and systemic *M. luteus* infection (P= 0.9999) (**Fig. 4D, Table S4, S5)**. At 12h and 48h, *M. luteus* colonies persisted in the heads of *Dif^1^* mutant post-brain (**Fig. 4C, Table S4, S5)** and systemic (**Fig. 4D, Table S4, S5)** infection. However, *M. luteus* was cleared in the head of control flies post-brain and systemic infection at respective time points. Together, these results indicate that host functional immunity clears the pathogen load, and long-term activated immunity is not associated with the presence of live bacteria in the brain.

### Glial cells primarily activate the brain’s immune response after E. coli and M. luteus brain infection

To determine whether glia, neurons or both are the primary source of AMP expression during the early and later stages of infection (24h and 2w-post-brain infection), we performed immunohistochemistry (IHC) on *Attacin::GFP* (*Att::GFP*), an AMP primarily regulated by the IMD pathway in response to Gram-negative bacteria, and *Drosomycin::GFP* (*Drs::GFP*) brains that express Repo in glial cells and Elav in neuronal cells. In *Att::GFP* flies, we observed that in response to sterile injury, some GFP expression was seen to co-localize with glia and neuronal markers at both 24h and 2 weeks post injury **(Fig. 5A, B, C)**; however, it was to a much higher extent in the *E. coli*-infected brains at both timepoints **(Fig. 5A, B, C).** These results suggest that glia are the primary cell type in which the IMD pathway is acutely and persistently activated after *E.coli* brain infection. We performed IHC in a second reporter line, *Cecropin::GFP*, for which we also found that *E. coli* brain infection resulted in upregulation of GFP at 24 hpi, which co-localized with the glial marker Repo **(Fig. S3 A, B).** While some neuronal expression of GFP is observed, these results support the idea that glia are the main cell type in which the IMD pathway is activated after *E. coli* infection.

**Figure 4.**
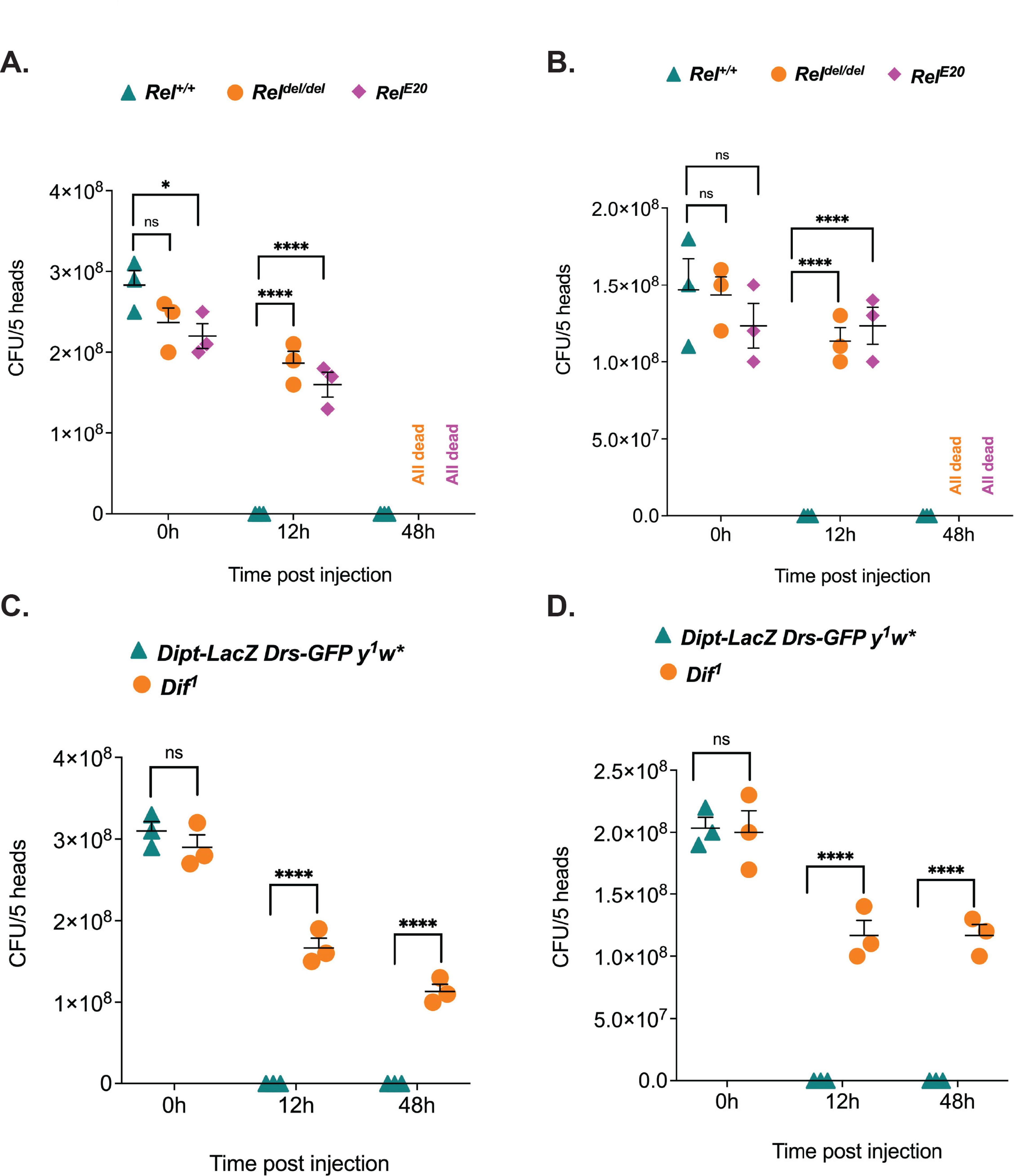
Functional immunity clears bacterial load in the *Drosophila* heads post-brain and systemic infection. **(A)** Ampicillin-resistant *E. coli* colonies were counted in *Rel^+/+^* (*Rel^del^-*control) and two *Relish* mutants (*Rel^del^* mutant and *Rel^E20^*) heads post-brain infection and **(B)** post-systemic infection at different time points. **(C)** *M. luteus* colonies were counted in control *Dipt-LacZ, Drs-GFP y^1^w\** and *Dif^1^*mutant post-brain infection and **(D)** post-systemic infection at different time points. Bacterial counts were done by plating the adult homogenates of 5 *Drosophila* heads that were previously infected with ampicillin-resistant *E. coli* and *M. luteus* in the brain and the thorax (systemic) on TSA plates containing ampicillin. The data shown are the number of colony-forming units (CFU) per 5 *Drosophila* head/replicate obtained at various time points post-brain and systemic bacterial infection. In all experiments, colonies counted at 0h represent the immediate bacterial load without aging the flies’ post-infection. Mean ± SEM; ***P < .005 based on a 2-way ANOVA test, with Tukey’s post-test for multiple comparisons. ns, not significant. Asterisks shown above the standard deviation bars denote significant differences in the pairwise control comparisons.

**Figure 5.**
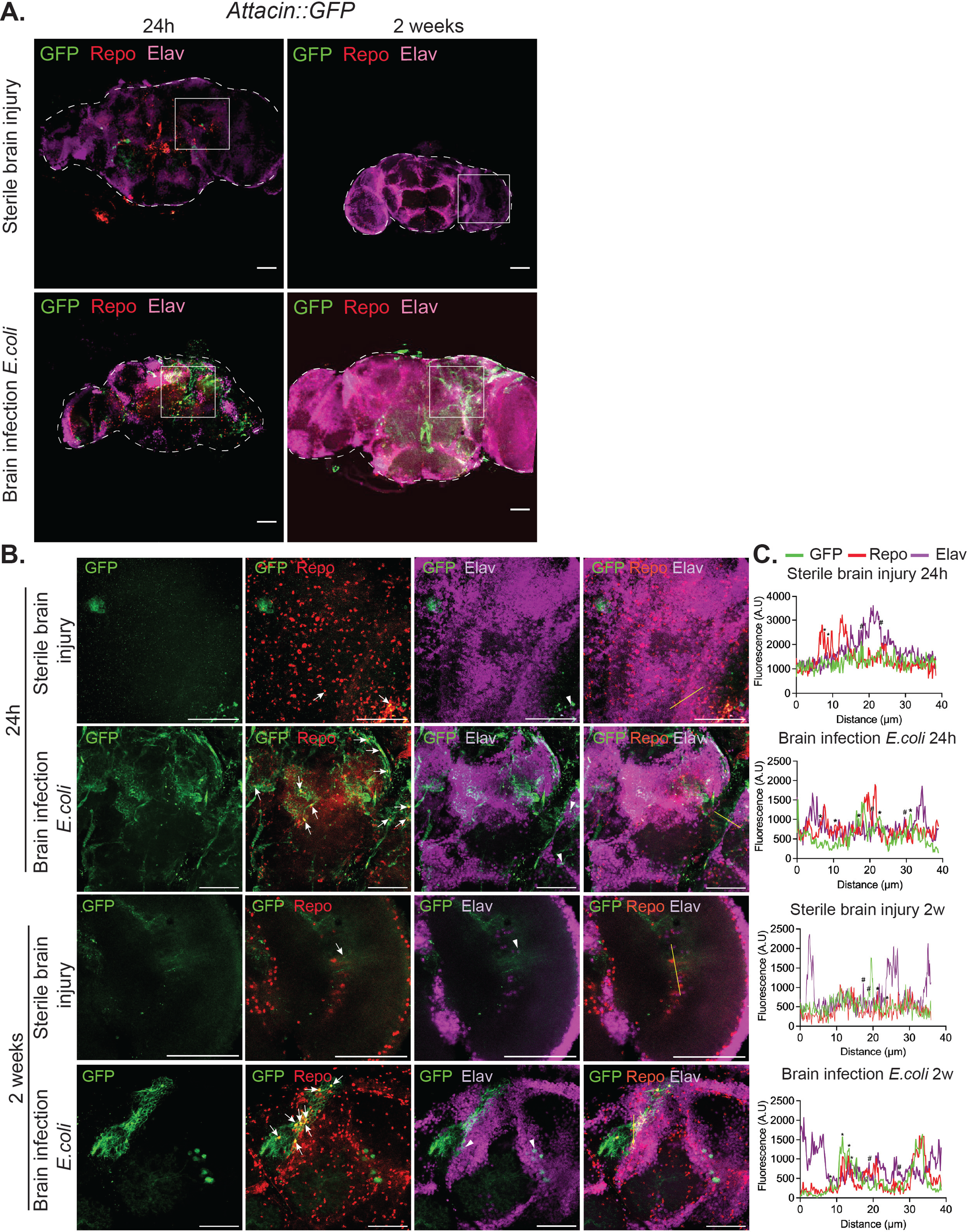
Glial cells are the primary source of acute and chronic AMP expression in *Attacin::GFP* brains post-*E. coli* brain infection. **(A)** Representative confocal images (20X) of 5-day-old *Att::GFP* brains 24h and 2w post-sterile injury and *E. coli* brain infection, immunostained for GFP (green), the glial marker Repo (red), and the neuronal marker Elav (magenta). *Drosophila* brain tissue is delineated with the discontinued line. A maximum projection of 3 z-stack slices per sample from n=3 brains were examined per experimental condition. White square boxes indicate the region of interest that was examined at higher magnification. **(B)** Representative confocal images (40X) of the area marked by the white square boxes in **(A)** of 5-day-old *Att::GFP* flies 24h and 2w post-sterile brain injury and *E. coli* brain infection. Co-localization between GFP and Repo is indicated with arrows, while co-localization between GFP and Elav is indicated with the arrowhead. Scalebars: 50µm. **(C)** Fluorescence intensity plots in selected brain areas (indicated by the yellow lines on the image merging GFP, Repo, and Elav) for the indicated samples. Overlapping peaks indicating co-localization between GFP and Repo (asterisks) and between GFP and Elav (pound symbol) are shown. A.U.: arbitrary units.

We next tested how *M. luteus* infection affected the acute and persistent expression of GFP in the brain of *Drs::GFP* flies, used as a readout for the Toll pathway. In *Drs::GFP* flies, we observed that in response to sterile injury, some GFP expressions were seen to co-localize with glia and neuronal markers at both 24h and 2 weeks post injury **(Fig. 6A, B, C)**; however, it was to a much higher extent in the *M. luteus*-infected brains at both timepoints **(Fig. 6A, B, C)**. The glial co-localization is more pronounced in the control-injected brain at 24h post-sterile brain injury compared to 2 weeks post-injection, suggesting an effect of the injury alone. In the case of *M. luteus* brain infection, more glial and less neuronal co-localization was observed at 2 weeks post-*M. luteus* infection, suggesting that the sustained inflammatory response showed continued glial activity and marked glial co-localization with the reporter. Altogether, these results indicate that glial cells are the primary cell type in the *Drosophila* brain that activate NF-kB immunity after acute and long-term bacterial brain infection.

**Figure 6.**
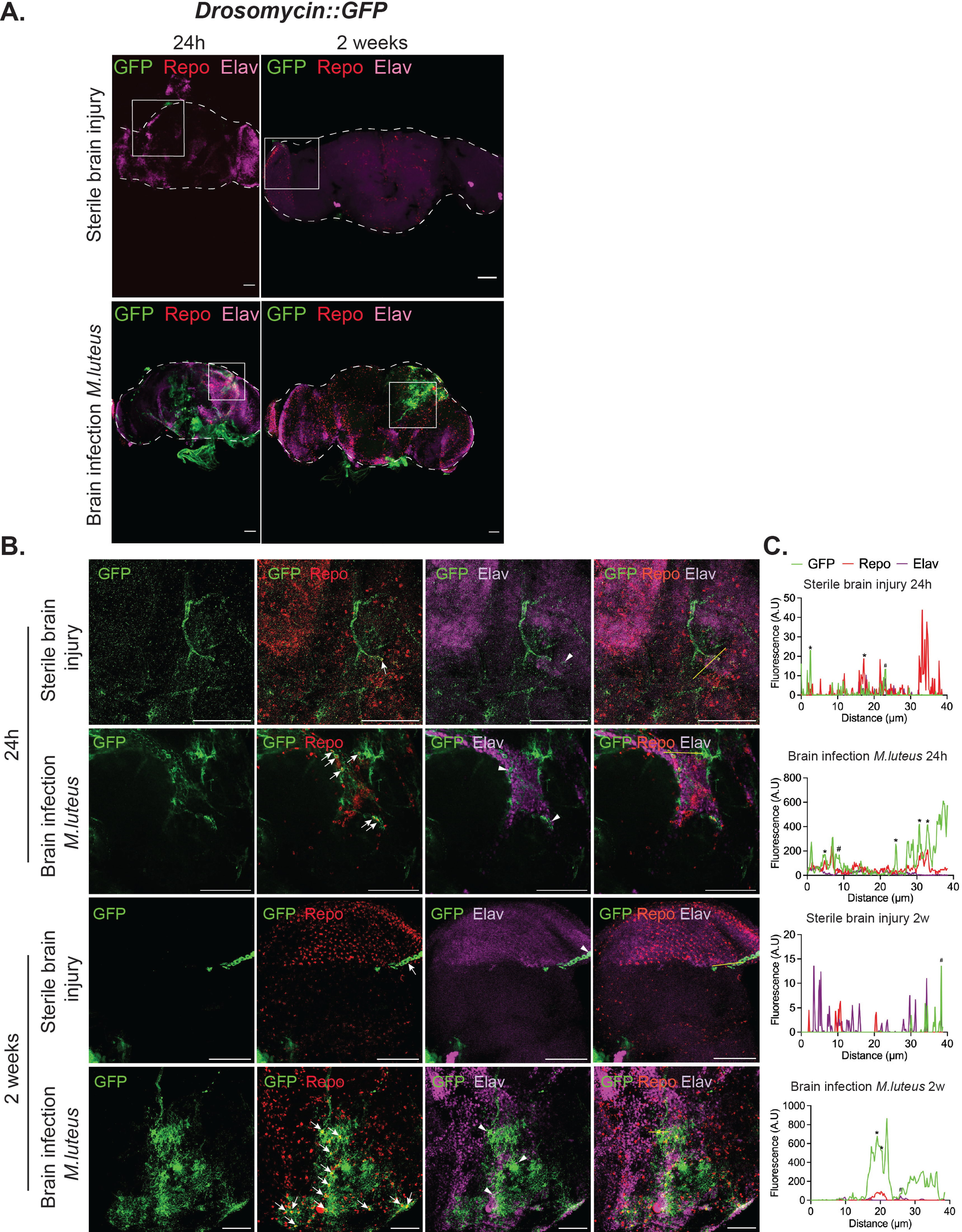
Glial cells are the primary source of acute and chronic AMP expression in *Drosomycin::GFP* brains post-*M. luteus* brain infection. **(A)** Representative confocal images (20X) of 5-day-old*, Drs::GFP* brains 24h and 2w post-sterile injury and *M. luteus* brain infection expressing reporter GFP in both glia and neurons and immunostained for GFP (green), the glial marker Repo (red), and the neuronal marker Elav (magenta). *Drosophila* brain tissue is delineated with the discontinued line. A maximum projection of 3 z-stack slices per sample from n=3 brains were examined per experimental condition. White square boxes indicate the region of interest that was examined at higher magnification. **(B)** Representative confocal images (40X) of the area marked by the white square boxes in **(A)** of 5-day-old *Drs::GFP* flies 24h and 2w post-sterile brain injury, and *M. luteus* brain infection. Co-localization between GFP and Repo is indicated with arrows, while co-localization between GFP and Elav is indicated with the arrowhead. Scalebars: 50µm. **(C)** Fluorescence intensity plots in selected brain areas (indicated by the yellow lines on the image merging GFP, Repo, and Elav) for the indicated samples. Overlapping peaks indicating co-localization between GFP and Repo (asterisks) and between GFP and Elav (pound symbol) are shown. A.U.: arbitrary units.

## Discussion

This study shows that AMP expression is activated acutely and chronically in the brain by both brain-specific and systemic bacterial infection. To understand the underlying mechanism, we used Toll and IMD pathway mutants and compared immune gene expression to their wild-type counterparts. Through this approach, we assessed the *Drosomycin* (*Drs*) and *Diptericin B* (*DiptB*) gene expression in the *Drosophila* brain to precisely determine the pathways’ involvement in activating brain innate immunity under various treatment conditions. *Tak1* and *Dredd* are crucial components of the IMD pathway that activate NF-κB signaling in the *Drosophila* brain following an *E. coli* infection (Leulier et al., 2000; Vidal et al., 2001). We directly compared *DiptB* expression in *y^1^w^67c23^* (*wild-type*) vs mutants (*Tak1* and *Dredd)* brains in the case of brain *E. coli* infection and systemic *E. coli* infection. Previous work has shown that *Tak1* and *Dredd* mutants are highly susceptible to gram-negative bacterial infection, failing to upregulate the *Diptericin* gene after *E. coli* infection (Leulier et al., 2000; Vidal et al., 2001). Consistent with this finding, we observed that both IMD pathway mutants showed impaired *DiptB* induction in the brain compared to *wild type*, following brain and systemic infection. Furthermore, in *y^1^w^67c23^* controls, we noted that direct *E. coli* brain infection showed higher *DiptB* expression in the brain compared to systemic *E. coli* infection. These results highlight that brain-specific *DiptB* expression is IMD pathway-dependent and validate the function of the IMD pathway components in controlling and mediating the brain’s innate immune response.

Our results showed the elevated *DiptB* expression in the brain 6h post-*E*. *coli* brain infection compared to non-injected (P<0.0001) and systemic infection (P=0.0003) **(Table S3)**. We observed *DiptB* activation at 12h, 24h, 36h, 48h, and up to 2 weeks post-brain infection, indicating the NF-κB activation profile ranging from acute to chronic. While the IMD pathway is highly effective against fast-replicating bacteria like *E. coli* because the pathway response via *DiptB* activation is quick ∼6 hpi (Buchon et al., 2009; Lemaitre and Hoffmann, 2007; Lemaitre et al., 1997) we assessed if activated immunity would clear the *E. coli* load in the brain over time. We decided to count *E. coli* load in the brain post-direct brain infection and systemic infection; therefore, *Rel*^+/+^ and two *Relish* mutants (*Rel^del/del^* and *Rel^E20^*) were infected with Ampicillin-resistant *E. coli* (HB101 + pGLO) directly in the head and on the thorax. The ampicillin-resistant *E. coli* was used to determine if the bacteria are cleared or if they continue to replicate. If this was the case, consequently, this could result in a long-term activation of immunity in the brain is due to the persistence of bacteria. We used *Rel* mutants as a positive control in our CFU assay, in which *E. coli* proliferated at 12h post-brain and systemic infection, and both mutants tested died at 48h post-brain and systemic infection. In contrast, in the heads of *Rel^+/+^* (controls), the *E. coli* load at 12h post-brain and systemic infections significantly dropped to zero colonies, indicating complete bacterial clearance in the head. This suggests that the functional IMD pathway is necessary to clear *E.coli* from the head after infection.

The dynamics of host resistance might change throughout the course of infection (Howick and Lazzaro, 2017). A previous study that injected 18.4nL of *E. coli* in the thorax measured bacterial load as CFU per fly and showed that *E. coli* load significantly reduced over time, with zero mortality at 72h post-infection, considering 24h and 72h as acute infection phase time-points and 1 week as a long-term chronic infection phase in their experiment. Another study (Kutzer and Armitage, 2016) showed that the host didn’t clear the *E. coli* load at 1-week p.i. However, our results showed complete *E. coli* clearance in the head at 12h post brain infection, as well as in a whole fly 12h post systemic infection **(Fig. S2, Table S4)**. We believe that *E. coli* infection in the brain and thorax activated the IMD pathway and had been effectively kept in check by the host immune system, failing to proliferate in number. It has been reported that activated antimicrobial defenses can act in the host hemolymph for weeks post-infection, most probably to control persistent bacterial infections (Makarova et al., 2016a, b). Nevertheless, various factors such as *E. coli* strain, sex, mating status, treatment (feeding vs non-feeding), and mode of bacterial introduction (pinprick vs dose nanoinjector) could largely contribute to discrepancies in the *E. coli* load over time (Khalil et al., 2015).

We wanted to further determine if *DiptB* expression in the brain resolved after bacterial clearance or persisted over time. Despite bacterial clearance, we found that *DiptB* is significantly induced in the brain up to 2 weeks, indicating chronic immune activation. The acute short-term activated IMD pathway in the context of brain injury is crucial for survival of infection and tissue repair (Nayak and Mishra, 2022; Zhai et al., 2018) whereas several studies have reported that prolonged activation of IMD signaling is detrimental because it promotes neuroinflammation, causing neurodegeneration (Kounatidis and Chtarbanova, 2018). In both vertebrates and invertebrates, chronic activation of glial cells is a hallmark of neuroinflammation (Garschall and Flatt, 2018; Kounatidis and Chtarbanova, 2018; Petersen et al., 2013). The study led by (Cao et al., 2013) used a mixture of *E. coli* and *M. luteus* to assess if an activated immune response in the CNS triggers neurodegeneration. They showed that persistently activated AMP expressions in glia or neurons can trigger neurodegeneration. Here, we utilized bacteria (*E. coli* and *M. luteus*) separately in our experiments to infect the *Drosophila* brain directly in the head and in the thorax. In agreement with the findings of Cao *et al*., our RT-qPCR results show elevated *DiptB* expression in the *Drosophila* brain. Additionally, immunostaining showed continued glial and some neuronal co-localization of *GFP* (for the IMD pathway reporter *Att::GFP*) in direct *E. coli*-infected brain at early (24h) and later (2 weeks) stages of infection. These sustained glial-mediated immune responses for an extended period indicate the shift from a transient acute response to a chronic one, raising the possibility of subsequent neuronal damage. In concurrence with the RT-qPCR results, our immunostaining results detected the reporter GFP signal in dissected brains when the injection was done in the head. Our results also align with previous findings showing that *Att::GFP* reporter expression in the brain was activated following brain-specific, but not systemic injection of a mixture of *E.coli* and *M.luteus*. Additionally, a lower GFP signal in control flies (sterile-injured flies), further confirms preferential activation of the immune response in the brain in bacterially injected flies. Therefore, the canonical IMD pathway mediates *DiptB* gene expression in the brain following bacterial brain infections, and systemic infection in the thorax doesn’t contribute to affecting gene expression in the brain over time.

The Toll pathway is another key regulator of NF-κB signaling in *Drosophila* that primarily defends against Gram-positive bacteria and fungi (Lemaitre and Hoffmann, 2007). MyD88 and Dif are crucial components of the Toll pathway, where *Drosophila* MyD88 (dMyD88) is an adaptor protein playing a similar role for inflammatory signaling pathways downstream of mammalian Toll-like receptor (TLR) (Horng and Medzhitov, 2001) while Dif is the main transcription factor acting downstream of the Toll receptor, mediating innate immune response via AMPs, including *Drs* gene expression (Lemaitre and Hoffmann, 2007). We directly compared the *Drs* expression in *w^1118^* and *Dipt-LacZ, Drs-GFP wild-type* controls vs *Myd88* and *Dif^1^* mutant brains in the case of both brain and systemic *M. luteus* infection. *Bomanin* genes are considered another set of Toll pathway-specific targets (Clemmons et al., 2015). Therefore, we measured *Bomanin* gene as an additional readout of the Toll pathway. We observed that *BomS2* gene expression was reduced in Toll pathway mutant’s brain compared to their controls at 24h post brain and systemic *M.luteus* infection **(Fig. S1) (Table S1)**. Our results showed that both *MyD88* and *Dif^1^* mutants failed to induce *Drs* gene expression in the brain upon direct brain and systemic *M. luteus* infection. However, interestingly, we observed that *Drs* gene was significantly induced in the wild-type brain post-systemic *M. luteus* infection. We didn’t expect bacterial infection in the thorax to induce an immune response in the brain, nor AMPs expressed in the fat body or in the cells outside of the CNS to enter the brain. Several studies have highlighted the significant evidence of neurodegenerative disease, not only as an outcome of CNS disorders, but also because of inter-organ communication. For instance, in a *Drosophila* model of Alzheimer’s disease (AD), enteric infection increased hemocyte motility and recruitment to the brain with an increase in oxidative stress, highlighting the gut-brain crosstalk (Wu et al., 2017). In a fly model of Parkinson’s Disease (PD), systemic infection contributed to disease pathogenesis via an inflammatory cascade and its downstream effects on dopaminergic neurons (Nayak and Mishra, 2022). Following systemic infection, the *Drosophila* fat body, a crucial immune organ and the primary site of antimicrobial peptide (AMP) production, releases signaling molecules, cytokines, and various AMPs into the hemolymph. These secreted molecules travel through hemolymph and signal to the brain, influencing CNS immune responses (Wang et al., 2024; Yu et al., 2022). It has been reported that a leukocytic antimicrobial peptide, Bactenecin, which is mainly present in bovine neutrophils, can directly cause cytotoxicity to cultured neurons and glia (Radermacher et al., 1993). However, in the *Drosophila* model, there is no direct evidence of fat body-derived AMPs crossing the BBB and mounting an immune response in the brain upon systemic bacterial infection. However, following systemic infection, AMP expression has been observed in the *Drosophila* head, possibly due to a local or peripheral immune response (Lee et al., 2024). Based on our results, *Drs* gene expressed in the brain after systemic *M. luteus* infection could be due to signals from organs like the fat body and hemocytes activating the Toll pathway in the brain. They revealed that the systemic *M. luteus* infection upregulated *Drs* expression in the brain, and later, *Drs* induction was significantly suppressed in the brain when *Spz* was knocked down in the fat body (Vincent et al., 2022). Other research evidence in *Drosophila* larvae showed that Spatzle secreted from hemocytes acts as a signal to the fat body in activating the Toll pathway, leading to *Drs* expression. These findings demonstrate the critical role of the fat body in immune signaling to the brain and show a link between the cellular and humoral immune responses. This emphasizes the importance of inter-organ communication in the *Drosophila* innate immune response by orchestrating a cytokine-based regulatory signal that is analogous to the mammalian immune system (Shia et al., 2009).

Like for the IMD pathway, we observed persistent *Drs* activation in the brain of *y^1^ w^67C23^* (control strains for Toll pathway’s mutant) flies at 12h, 24h, 36h, 48h, and up to 2 weeks post-brain infection, indicating the NF-kB activation profile ranging from acute to chronic. At 12h post-brain infection, *M. luteus* load reduced to 0 colonies in *Dipt-LacZ, Drs-GFP y^1^w\** control brain, indicating complete bacterial clearance. Similarly, upon systemic infection, *M. luteus* load reduced to 0 colonies, suggesting that the activated Toll pathway contributed to host resistance via functional *Drs* activation. At both direct brain and systemic infection, *M. luteus* proliferated in the *Dif^1^*mutant’s head up to 48 hours, but the mutants didn’t die consistent with previous studies that this bacterium induces Toll-dependent *Drs* expression but doesn’t kill Toll pathway mutants but (De Gregorio et al., 2002; Rutschmann et al., 2000). Despite bacterial clearance, our RT-qPCR results showed elevated *Drs* expression in the brain for up to 2 weeks, coupled with immunostaining showing continued glial-specific co-localization of *Drs::GFP* (for *Drosomycin*) in direct *M. luteus-*infected brain at early (24h) and later (2 weeks) stages of infection. The RT-qPCR results showing induced *Drs* expression in the brain post-systemic *M. luteus* infection were further validated by staining results that showed mostly glial and some neuronal co-localization of *Drs::GFP* at 24 hpi **(Fig. S3 C, D, E) (Table S1)**. Previous studies have shown that heat-killed bacteria, similar to live bacteria, can activate the immune system and induce AMPs (Noh et al., 2022). It has been reported that oral administration of non-pathogenic gram-positive and gram-negative heat-killed bacteria activated Toll and IMD pathways, inducing *Drosomycin* and *Diptericin* genes, respectively (Wen et al., 2019). Concurrent to their observation of killed bacteria, our pin-prick method of heat-killed bacterial infection showed enhanced *DiptB* and *Drs* expression in the brain post-brain infection in both pathways. In the context of a diseased brain, such as penetrating traumatic brain injury (pTBI), there is evidence that the same AMPs were upregulated by both pathways (Marischuk et al., 2021). To understand the complexity of pathway activation in the context of bacterial brain infection, we decided to see if a crosstalk between the Toll and IMD NF-κB pathways exists, or if their activation in the brain happens independently. Thus, we measured both *DiptB* and *Drs* expression in the wild type control brain at 24h post *E.coli* infection **(Fig. S4 A) (Table S1)** and at 6h post *M.luteus* infection **(Fig. S4 B) (Table S1)** and observed that both *DiptB* and *Drs* genes were coregulated by both pathways, highlighting a possible crosstalk. A recent study highlighting the communication pathway between the brain and muscles during infection has revealed that acute CNS infections initiate a long-term reduction in motor function, which indicates that a systemic, rather than a direct neuronal mechanism, is at play (Yang et al., 2024). Our results showing long-term activated immunity in the brain, either via direct brain infection or systemic infection, could potentially exacerbate neuronal death through local inflammation, contributing to neurodegeneration (Lee et al., 2024; Stuart et al., 2022).

## Conclusion and limitations

We performed RT-qPCR on dissected brain post-brain and systemic bacterial infection and validated our immune gene expression results via immunostaining with specific tissue markers showing glia as the primary cell type activating immune response at early and later stages of infection. However, there are still some limitations to our study that warrant further consideration. To address the central question of whether long-term activated immune response is deleterious to neuronal health, we will require histological analysis to observe neuronal integrity and behavioral assay as a sensitive readout of neuronal dysfunction. Measuring the life span of those flies that cleared the bacteria after 12 hpi compared to the non-infected cohort would give us an idea about the overall fitness cost of chronic immune activation. To better understand the behavior of *Drosophila* post-infection, their locomotory activity could be measured via a climbing assay or an automated video tracking assay to see if neuroinflammation leads to either hyperactivity or causes a decline in their general activity (Bolus et al., 2020; Mainali et al., 2025). To further assess the cellular aspect of immune response, a phagocytic assay needs to be performed to see if the activated glia engulf the bacteria or if glial phagocytic ability is altered during the long-term immune activation that persists. We can extend the immunohistochemical analysis to other AMPs to better characterize glial cell subtypes involved in immune activation at early and later stages of infection. Our approach of using *Drosophila* as a model to study infection-induced neuroinflammation and its link to neuropathology aims to understand the role of various players in NF-κB pathways and characterize cell-type-specific mechanisms at both early and later stages of infection. This study advances knowledge of how non-lethal brain bacterial infection results in the activation of NF-κB immunity that could be targeted to prevent or halt progression of neurodegeneration.

## Acknowledgments

We are grateful to Dr. Kim Lackey and the Optical Analysis Facility for assistance with confocal microscopy, and to Patrice Crawford for kindly providing the bacterial culture used throughout our infection experiments. We would also like to thank Brantley Johnson for assistance with fly handling, amplifying, and maintenance of stocks. We also acknowledge the valuable intellectual input and helpful suggestions from our colleagues of the Ganetzky joint lab meeting.

## Authors contributions

SC conceptualized the study, with input from SM. SM, IT, PM, JG, LH, EK, KD and SC performed experiments; SM, IT, PM, JG, LH and SC analyzed data. SM wrote the original draft. SC contributed to manuscript writing, review and editing.

## Supplemental figures legends

**Supplemental Figure S1.**
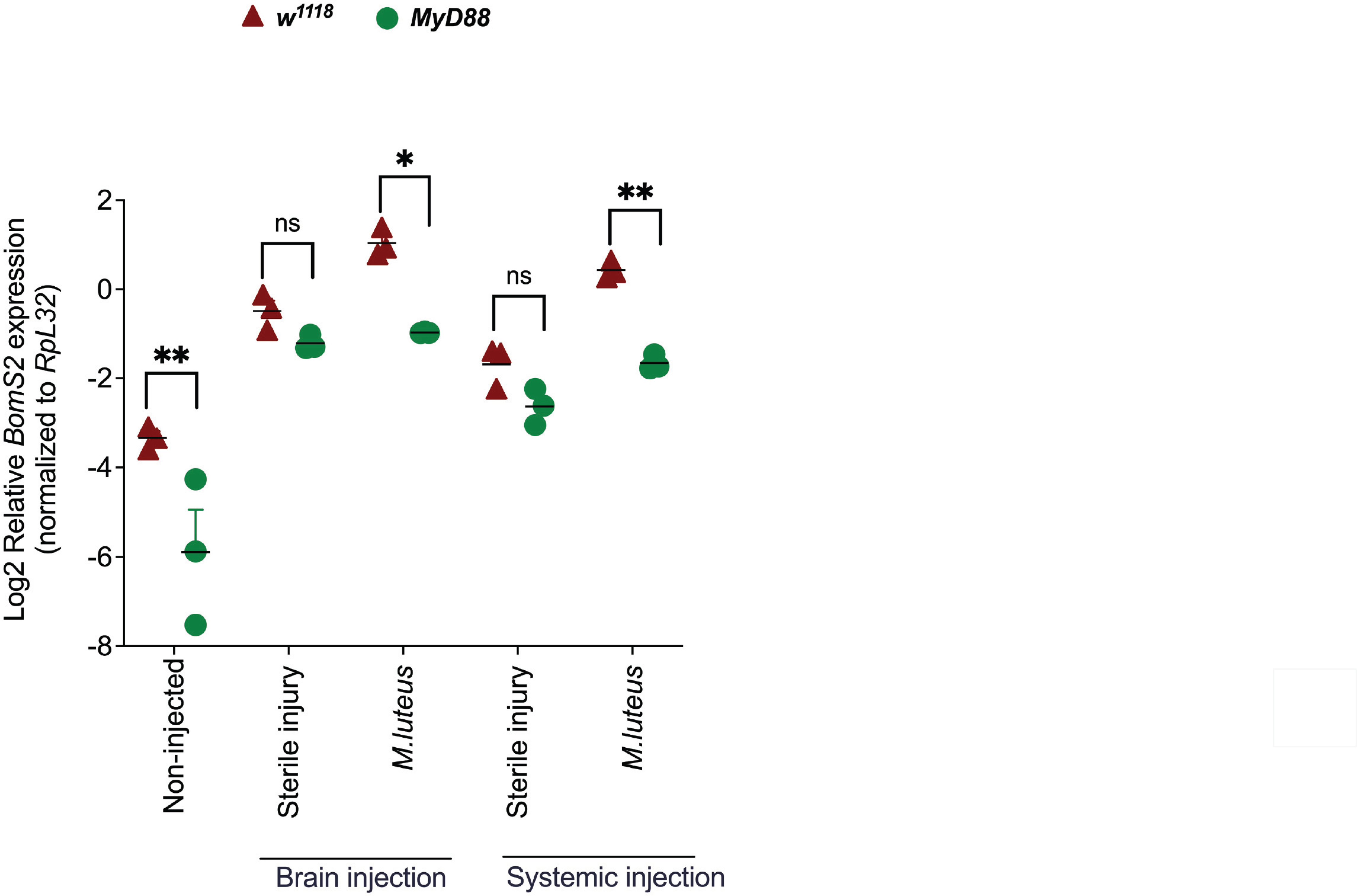
*BomS2* gene expression measured in controls and Toll pathway mutants’ brain tissue 24h post-head (brain) and thorax (systemic) *M. luteus* infection. N =15 dissected brains per replicate, with n=3 biological replicates (individual symbols on the graph) per treatment. *BomS2* expression levels were normalized to the housekeeping gene *RpL32* and log transformed. Mean ± SEM; ***P < .005 based on a 2-way ANOVA test, with Tukey’s post-test for multiple comparisons. ns, not significant. Asterisks shown above the standard deviation bars denote significant differences in the pairwise control comparisons.

**Supplemental Figure S2.**
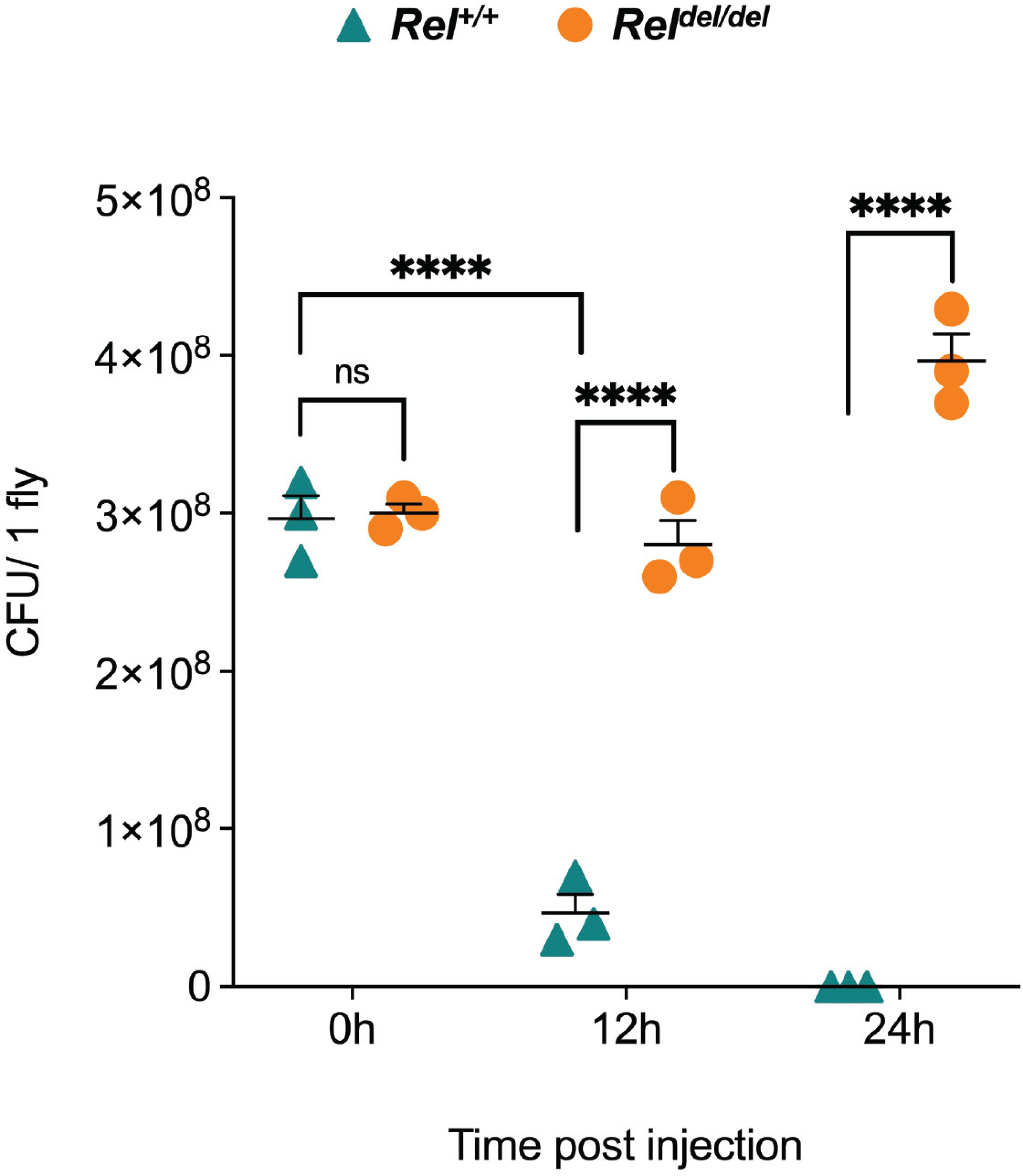
Ampicillin-resistant *E. coli* colonies were counted in the whole fly of *Rel^+/+^* (control) and *Rel^del/del^* mutants post-*E. coli* systemic infection at different time points. Bacterial counts were done by plating the homogenates of 1 whole *Drosophila* that were previously infected with ampicillin-resistant *E. coli* in the thorax (systemic) on TSA plates containing ampicillin. The data shown is the number of colony-forming units (CFU) per 1 *Drosophila*/replicate obtained at various time points post-systemic *E. coli* infection. Colonies counted at 0h represent the immediate bacterial load following *E. coli* introduction in the fly hemocoel. Mean ± SEM; ***P < .005 based on a 2-way ANOVA test, with Tukey’s post-test for multiple comparisons. ns, not significant. Asterisks shown above the standard deviation bars denote significant differences in the pairwise control comparisons.

**Supplemental Figure S3.**
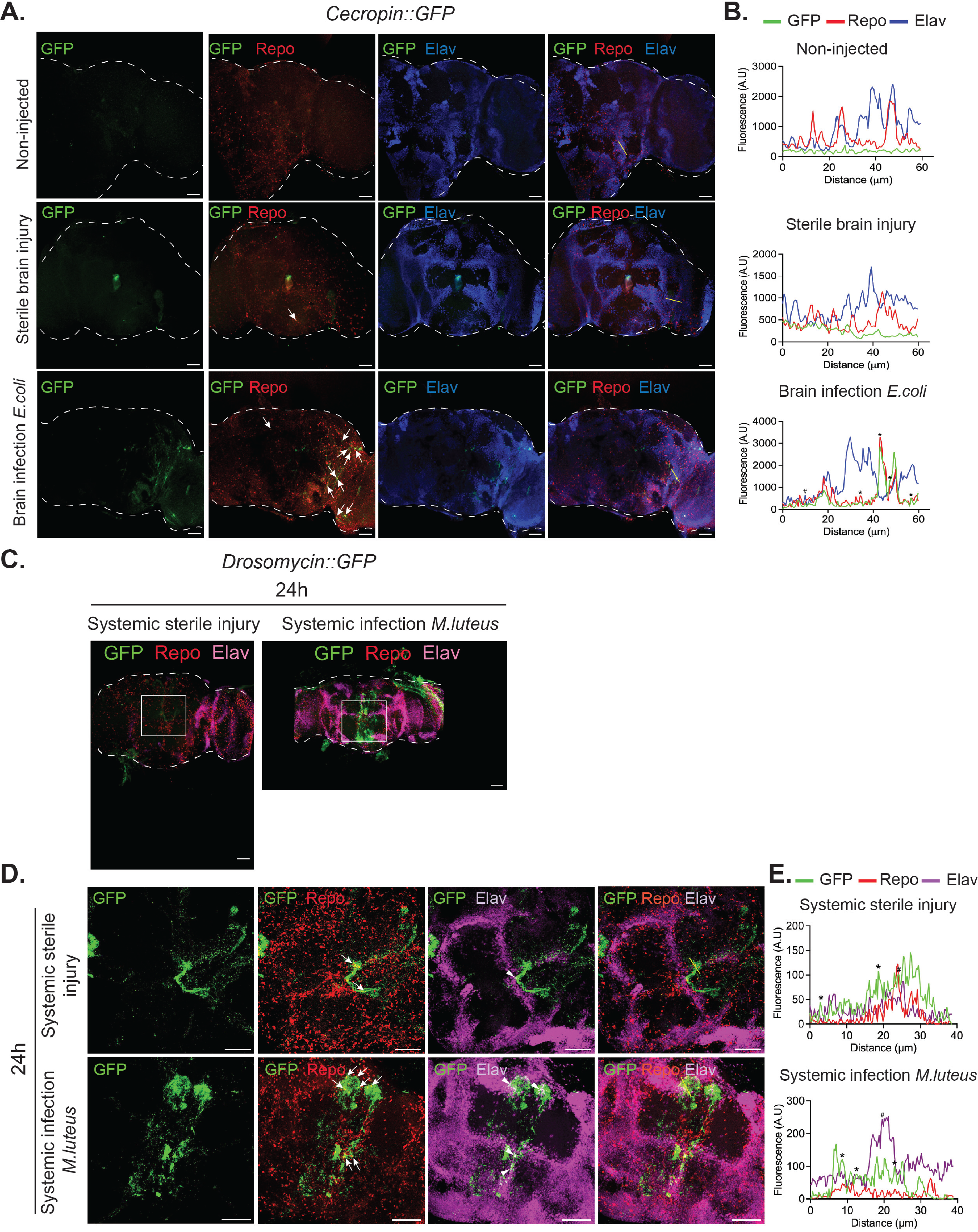
**(A)** Representative confocal stack images (20X) of 4-7 days old*, Cecropin::GFP* brains 24h post-*E. coli* brain infection. Brains of non-injected flies or sterile-injured brains were used as controls. *Drosophila* brain tissue is delineated with the discontinued line. Maximum projection of 5 slices per sample from n=3 brains was examined per experimental condition. Immunostaining for GFP (green), the glial marker Repo (red), and the neuronal marker Elav (blue) shows co-localization primarily between GFP-expressing cells and Repo (arrows) in *E. coli*-infected brains. **(B)** Fluorescence intensity plots in selected brain areas (indicated by the yellow lines on the image merging GFP, Repo, and Elav) for the indicated samples. Overlapping peaks indicating co-localization between GFP and Repo (asterisks) and between GFP and Elav (pound symbol) are shown. A.U.: arbitrary units. **(C)** Representative confocal images (20X) of 5-day-old*, Drs::GFP* brains 24h post-sterile thorax injury and *M. luteus* thorax infection (systemic infection), immunostained for GFP (green), the glial marker Repo (red), and the neuronal marker Elav (magenta). *Drosophila* brain tissue is delineated with the discontinued line. White square boxes indicate the region of interest that was examined at higher magnification. **(D)** Representative confocal images (40X) of 5-day-old *Drs::GFP*, GFP was colocalized with Repo in the brain 24h post systemic *M. luteus* infection, indicated by white arrow heads. Co-localization between GFP and Repo is indicated with arrows, while co-localization between GFP and Elav is indicated with the arrowhead. **(A, C, D)** Scalebars: 50 µm. **(E)** Fluorescence intensity plots in selected brain areas (indicated by the yellow lines on the image merging GFP, Repo, and Elav) for the indicated samples. Overlapping peaks indicating co-localization between GFP and Repo (asterisks) and between GFP and Elav (pound symbol) are shown. A.U.: arbitrary units.

**Supplemental Figure S4.**
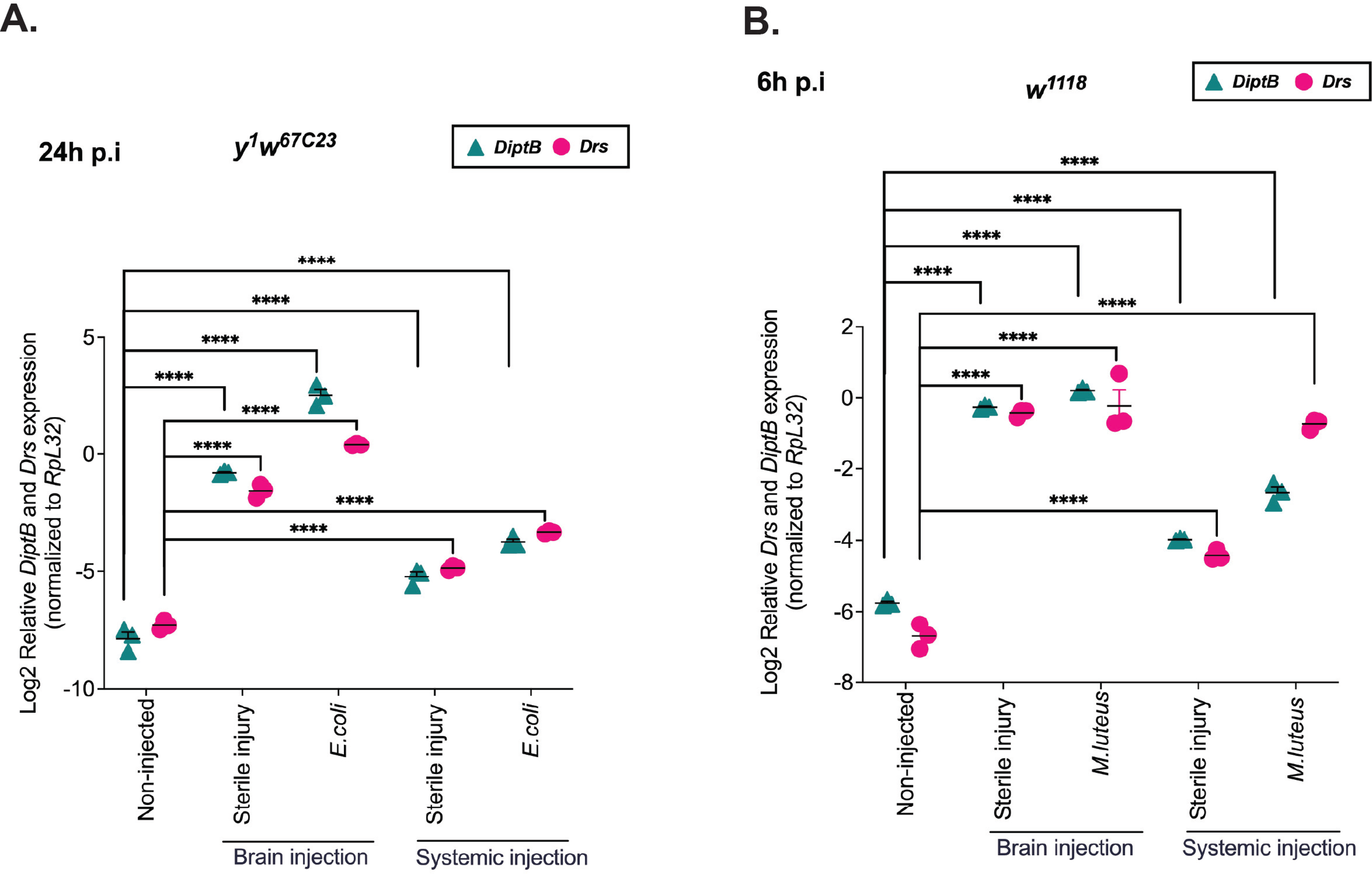
**(A)** *DiptB* and *Drs* gene expression were measured in the control *y^1^ w^67c23^* genotype brains 24h post-head (brain) and thorax (systemic) *E. coli* infection. N =15 dissected brains per replicate, with n=3 biological replicates (individual symbols on the graph) per treatment. *DiptB* and *Drs* expression levels were normalized to the housekeeping gene *RpL32* and log-transformed. Mean ± SEM; ***P < .005 based on a 2-way ANOVA test, with Tukey’s post-test for multiple comparisons. ns, not significant. Asterisks shown above the standard deviation bars denote significant differences in the pairwise control comparisons. **(B)** *Drs* and *DiptB* gene expression were measured in control *w^1118^* genotype brains 6h post-head (brain) and thorax (systemic) *M. luteus* infection. N =15 dissected brains were used per replicate, with n=3 biological replicates (individual symbols on the graph) per treatment. *Drs* and *DiptB* expression levels were normalized to the housekeeping gene *RpL32* and log transformed. Mean ± SEM; ***P < .005 based on a 2-way ANOVA test, with Tukey’s post-test for multiple comparisons. ns, not significant. Asterisks shown above the standard deviation bars denote significant differences in the pairwise control comparisons.

## References

Bolus, H., Crocker, K., Boekhoff-Falk, G., Chtarbanova, S., 2020. Modeling Neurodegenerative Disorders in Drosophila melanogaster. Int J Mol Sci 21.

Buchon, N., Broderick, N.A., Poidevin, M., Pradervand, S., Lemaitre, B., 2009. Drosophila intestinal response to bacterial infection: activation of host defense and stem cell proliferation. Cell Host Microbe 5, 200–211.

Cao, Y., Chtarbanova, S., Petersen, A.J., Ganetzky, B., 2013. Dnr1 mutations cause neurodegeneration in Drosophila by activating the innate immune response in the brain. Proc Natl Acad Sci U S A 110, E1752–1760.

Clemmons, A.W., Lindsay, S.A., Wasserman, S.A., 2015. An effector Peptide family required for Drosophila toll-mediated immunity. PLoS Pathog 11, e1004876.

De Gregorio, E., Spellman, P.T., Tzou, P., Rubin, G.M., Lemaitre, B., 2002. The Toll and Imd pathways are the major regulators of the immune response in Drosophila. Embo j 21, 2568–2579.

Garschall, K., Flatt, T., 2018. The interplay between immunity and aging in Drosophila. F1000Res 7, 160.

Horng, T., Medzhitov, R., 2001. Drosophila MyD88 is an adapter in the Toll signaling pathway. Proc Natl Acad Sci U S A 98, 12654–12658.

Howick, V.M., Lazzaro, B.P., 2017. The genetic architecture of defence as resistance to and tolerance of bacterial infection in Drosophila melanogaster. Mol Ecol 26, 1533–1546.

Jang, I.H., Chosa, N., Kim, S.H., Nam, H.J., Lemaitre, B., Ochiai, M., Kambris, Z., Brun, S., Hashimoto, C., Ashida, M., Brey, P.T., Lee, W.J., 2006. A Spatzle-processing enzyme required for toll signaling activation in Drosophila innate immunity. Dev Cell 10, 45–55.

Khalil, S., Jacobson, E., Chambers, M.C., Lazzaro, B.P., 2015. Systemic bacterial infection and immune defense phenotypes in Drosophila melanogaster. J Vis Exp, e52613.

Kounatidis, I., Chtarbanova, S., 2018. Role of Glial Immunity in Lifespan Determination: A Drosophila Perspective. Front Immunol 9, 1362.

Kounatidis, I., Chtarbanova, S., Cao, Y., Hayne, M., Jayanth, D., Ganetzky, B., Ligoxygakis, P., 2017. NF-kappaB Immunity in the Brain Determines Fly Lifespan in Healthy Aging and Age-Related Neurodegeneration. Cell Rep 19, 836–848.

Kutzer, M.A., Armitage, S.A., 2016. The effect of diet and time after bacterial infection on fecundity, resistance, and tolerance in Drosophila melanogaster. Ecol Evol 6, 4229–4242.

Larrea, A., Elexpe, A., Diez-Martin, E., Torrecilla, M., Astigarraga, E., Barreda-Gomez, G., 2023. Neuroinflammation in the Evolution of Motor Function in Stroke and Trauma Patients: Treatment and Potential Biomarkers. Curr Issues Mol Biol 45, 8552-8585.

Leclerc, V., Reichhart, J.M., 2004. The immune response of Drosophila melanogaster. Immunol Rev 198, 59–71.

Lee, K.Z., Ferrandon, D., 2011. Negative regulation of immune responses on the fly. Embo j 30, 988–990.

Lee, S., Silverman, N., Gao, F.B., 2024. Emerging roles of antimicrobial peptides in innate immunity, neuronal function, and neurodegeneration. Trends Neurosci 47, 949–961.

Lemaitre, B., Hoffmann, J., 2007. The host defense of Drosophila melanogaster. Annu Rev Immunol 25, 697–743.

Lemaitre, B., Reichhart, J.M., Hoffmann, J.A., 1997. Drosophila host defense: differential induction of antimicrobial peptide genes after infection by various classes of microorganisms. Proc Natl Acad Sci U S A 94, 14614–14619.

Leulier, F., Rodriguez, A., Khush, R.S., Abrams, J.M., Lemaitre, B., 2000. The Drosophila caspase Dredd is required to resist gram-negative bacterial infection. EMBO Rep 1, 353–358.

Liegeois, S., Ferrandon, D., 2022. Sensing microbial infections in the Drosophila melanogaster genetic model organism. Immunogenetics 74, 35–62.

Lotz, S.K., Blackhurst, B.M., Reagin, K.L., Funk, K.E., 2021. Microbial Infections Are a Risk Factor for Neurodegenerative Diseases. Front Cell Neurosci 15, 691136.

Lye, S.H., Chtarbanova, S., 2018. Drosophila as a Model to Study Brain Innate Immunity in Health and Disease. Int J Mol Sci 19.

Mainali, S., Majlish, A.N.K., Lee, Y.R., Lee, H., Iyengar, A., Chtarbanova, S., 2025. A Tool Kit to Model Neurodegenerative Disease in Drosophila melanogaster, in: Muñoz-Torrero, D. (Ed.), Methods in Neurodegenerative Disease Drug Discovery. Springer US, New York, NY, pp. 283–312.

Makarova, O., Rodriguez-Rojas, A., Eravci, M., Weise, C., Dobson, A., Johnston, P., Rolff, J., 2016a. Antimicrobial defence and persistent infection in insects revisited. Philos Trans R Soc Lond B Biol Sci 371.

Makarova, O., Rodriguez-Rojas, A., Eravci, M., Weise, C., Dobson, A., Johnston, P., Rolff, J., 2016b. Correction to: ‘Antimicrobial defence and persistent infection in insects revisited’. Philos Trans R Soc Lond B Biol Sci 371.

Marischuk, K., Crocker, K.L., Ahern-Djamali, S., Boekhoff-Falk, G., 2021. Innate immunity pathways activate cell proliferation after penetrating traumatic brain injury in adult *Drosophila*. bioRxiv, 2021.2009.2001.458615.

Myllymaki, H., Valanne, S., Ramet, M., 2014. The Drosophila imd signaling pathway. J Immunol 192, 3455–3462.

Nayak, N., Mishra, M., 2022. Drosophila melanogaster as a model to understand the mechanisms of infection mediated neuroinflammation in neurodegenerative diseases. J Integr Neurosci 21, 66.

Noh, H.J., Park, J.M., Kwon, Y.J., Kim, K., Park, S.Y., Kim, I., Lim, J.H., Kim, B.K., Kim, B.Y., 2022. Immunostimulatory Effect of Heat-Killed Probiotics on RAW264.7 Macrophages. J Microbiol Biotechnol 32, 638–644.

Petersen, A.J., Katzenberger, R.J., Wassarman, D.A., 2013. The innate immune response transcription factor relish is necessary for neurodegeneration in a Drosophila model of ataxia-telangiectasia. Genetics 194, 133–142.

Postolache, T.T., Wadhawan, A., Can, A., Lowry, C.A., Woodbury, M., Makkar, H., Hoisington, A.J., Scott, A.J., Potocki, E., Benros, M.E., Stiller, J.W., 2020. Inflammation in Traumatic Brain Injury. J Alzheimers Dis 74, 1–28.

Radermacher, S.W., Schoop, V.M., Schluesener, H.J., 1993. Bactenecin, a leukocytic antimicrobial peptide, is cytotoxic to neuronal and glial cells. J Neurosci Res 36, 657–662.

Rodríguez, A.M., Rodríguez, J., Giambartolomei, G.H., 2022. Microglia at the Crossroads of Pathogen-Induced Neuroinflammation. ASN Neuro 14, 17590914221104566.

Rutschmann, S., Jung, A.C., Hetru, C., Reichhart, J.M., Hoffmann, J.A., Ferrandon, D., 2000. The Rel protein DIF mediates the antifungal but not the antibacterial host defense in Drosophila. Immunity 12, 569–580.

Shan, T., Wang, Y., Bhattarai, K., Jiang, H., 2023. An evolutionarily conserved serine protease network mediates melanization and Toll activation in Drosophila. Sci Adv 9, eadk2756.

Shia, A.K., Glittenberg, M., Thompson, G., Weber, A.N., Reichhart, J.M., Ligoxygakis, P., 2009. Toll-dependent antimicrobial responses in Drosophila larval fat body require Spatzle secreted by haemocytes. J Cell Sci 122, 4505–4515.

Stuart, B.A.R., Franitza, A.L., E, L., 2022. Regulatory Roles of Antimicrobial Peptides in the Nervous System: Implications for Neuronal Aging. Front Cell Neurosci 16, 843790.

Swanson, L.C., Rimkus, S.A., Ganetzky, B., Wassarman, D.A., 2020a. Loss of the Antimicrobial Peptide Metchnikowin Protects Against Traumatic Brain Injury Outcomes in Drosophila melanogaster. G3 (Bethesda) 10, 3109–3119.

Swanson, L.C., Trujillo, E.A., Thiede, G.H., Katzenberger, R.J., Shishkova, E., Coon, J.J., Ganetzky, B., Wassarman, D.A., 2020b. Survival Following Traumatic Brain Injury in Drosophila Is Increased by Heterozygosity for a Mutation of the NF-κB Innate Immune Response Transcription Factor Relish. Genetics 216, 1117–1136.

Tauszig-Delamasure, S., Bilak, H., Capovilla, M., Hoffmann, J.A., Imler, J.L., 2002. Drosophila MyD88 is required for the response to fungal and Gram-positive bacterial infections. Nat Immunol 3, 91–97.

Tzou, P., Ohresser, S., Ferrandon, D., Capovilla, M., Reichhart, J.M., Lemaitre, B., Hoffmann, J.A., Imler, J.L., 2000. Tissue-specific inducible expression of antimicrobial peptide genes in Drosophila surface epithelia. Immunity 13, 737–748.

Vidal, S., Khush, R.S., Leulier, F., Tzou, P., Nakamura, M., Lemaitre, B., 2001. Mutations in the Drosophila dTAK1 gene reveal a conserved function for MAPKKKs in the control of rel/NF- kappaB-dependent innate immune responses. Genes Dev 15, 1900–1912.

Vincent, C.M., Beckwith, E.J., Simoes da Silva, C.J., Pearson, W.H., Kierdorf, K., Gilestro, G.F., Dionne, M.S., 2022. Infection increases activity via Toll dependent and independent mechanisms in Drosophila melanogaster. PLoS Pathog 18, e1010826.

Wang, Y., Saito, K., Tanimoto, H., Grunwald Kadow, I.C., 2024. A Bidirectional Brain-Fat Body Axis for Pathogen Avoidance. bioRxiv, 2024.2011.2002.621634.

Wen, Y., He, Z., Xu, T., Jiao, Y., Liu, X., Wang, Y.F., Yu, X.Q., 2019. Ingestion of killed bacteria activates antimicrobial peptide genes in Drosophila melanogaster and protects flies from septic infection. Dev Comp Immunol 95, 10–18.

Winkler, B., Funke, D., Benmimoun, B., Speder, P., Rey, S., Logan, M.A., Klambt, C., 2021. Brain inflammation triggers macrophage invasion across the blood-brain barrier in Drosophila during pupal stages. Sci Adv 7, eabh0050.

Winkler, B., Funke, D., Klambt, C., 2025. Macrophage invasion into the Drosophila brain requires JAK/STAT-dependent MMP activation in the blood-brain barrier. PLoS Biol 23, e3003035.

Wu, S.C., Cao, Z.S., Chang, K.M., Juang, J.L., 2017. Intestinal microbial dysbiosis aggravates the progression of Alzheimer’s disease in Drosophila. Nat Commun 8, 24.

Xu, R., Lou, Y., Tidu, A., Bulet, P., Heinekamp, T., Martin, F., Brakhage, A., Li, Z., Liégeois, S., Ferrandon, D., 2023. The Toll pathway mediates *Drosophila* resilience to *Aspergillus* mycotoxins through specific Bomanins. EMBO reports 24, e56036.

Yang, S., Tian, M., Dai, Y., Wang, R., Yamada, S., Feng, S., Wang, Y., Chhangani, D., Ou, T., Li, W., Guo, X., McAdow, J., Rincon-Limas, D.E., Yin, X., Tai, W., Cheng, G., Johnson, A., 2024. Infection and chronic disease activate a systemic brain-muscle signaling axis. Sci Immunol 9, eadm7908.

Yu, S., Luo, F., Xu, Y., Zhang, Y., Jin, L.H., 2022. Drosophila Innate Immunity Involves Multiple Signaling Pathways and Coordinated Communication Between Different Tissues. Front Immunol 13, 905370.

Zhai, Z., Huang, X., Yin, Y., 2018. Beyond immunity: The Imd pathway as a coordinator of host defense, organismal physiology and behavior. Dev Comp Immunol 83, 51–59.

Zindler, E., Zipp, F., 2010. Neuronal injury in chronic CNS inflammation. Best Pract Res Clin Anaesthesiol 24, 551–562.

